# Mechanisms of sex differences in acute and long COVID sequelae in mice

**DOI:** 10.1101/2025.10.13.682101

**Authors:** Jennifer A. Liu, Sabal Chaulagain, Xinyi Xie, Patrick S. Creisher, Weizhi Zhong, Tianle Zhang, Maraake Taddese, Karly Shi, Han-Sol Park, Haley Hcnir, Arthur P. Arnold, Ralph Baric, Natasha B. Barahona, Elizabeth Engler-Chiurazzi, Kevin J. Zwezdaryk, Chloe L. Thio, Ashwin Balagopal, Jack R. Harkema, Elizabeth A. Thompson, Andrew Pekosz, Andrea L. Cox, Sabra L. Klein

## Abstract

Males suffer more severe outcomes during acute COVID-19 and a greater proportion of females develop post-acute sequalae of COVID-19 (PASC). To identify mechanisms of PASC, mice were infected with SARS-CoV-2 and viral, inflammatory, and behavioral outcomes were evaluated through 84 days post-infection. Sex differences were not observed in virus replication or persistence of viral RNA in pulmonary or extrapulmonary tissues. After infectious virus clearance, females exhibited persistent deficits in spatial working and recognition memory, and greater frequencies of inflammatory myeloid cells, neuroinflammation, and dysregulated T cells, including Tregs. Sex differences in persistent inflammation and cognitive impairments during PASC were mediated by the presence of two X chromosomes and infection-induced upregulation of the X-linked genes *Xist* and *Tlr7*, which was reversed by pharmacological inhibition of TLR7 activity during the PASC phase. Sustained X-linked TLR7-mediated inflammation underlies how the burden of PASC is greater for females and provides a therapeutic target.

## Introduction

Approximately 5-10% of patients with COVID-19 experience prolonged symptoms and disease after SARS-CoV-2 infection, referred to as post-acute sequelae of COVID-19 (PASC) or long COVID^1^. Although SARS-CoV-2 is a respiratory pathogen that typically remains restricted to the lungs, PASC is a heterogenous disease with symptoms spanning across multi-organs, with neurological and cognitive symptoms comprising a significant proportion of clinical presentations of disease^2–5^. Global estimates project that 400 million people worldwide have experienced PASC^6^ including over 18 million in the U.S.^7^. Despite diagnostic definitions of PASC^1,8^, there are still no approved disease modifying therapeutic treatments, in part because the underlying basic mechanisms of disease remain elusive, and pathophysiological mechanisms likely differ between disease subtypes or other factors, including biological sex. It has been well established that sex differences are observed in acute COVID-19 and PASC outcomes.

Data from the U.S. National Institutes of Health Researching COVID to Enhance Recovery (RECOVER)^9^ adult cohort show that while males are more likely to be severely ill in acute COVID-19, females are more likely to develop PASC. While PASC occurs in all ages, across acute disease severities, and SARS-CoV-2 variants^1,10^, it predominantly occurs in middle aged populations following mild acute illness and in aged populations requiring hospitalization^1,11^ but the biological mechanisms underlying sex differences remain elusive in both the acute and PASC phases of SARS-CoV-2 infection.

Neurological features and cognitive symptoms, including impaired executive functioning, memory loss, “brain fog”, and fatigue are reported in approximately a third of individuals with PASC^12^, with neurocognitive features being more common in females than males independent of acute disease severity^2,5,13^. Despite the general absence of CNS infection outside of very severe COVID-19, long-term changes in the brain have been documented in patients with PASC, including reduced cortical thickness^14^, gray matter changes^15^, and blood-brain barrier disruption^16^, with more recent studies identifying distinct microstructural changes across multiple brain regions^3^ that vary by clinical symptom presentation (i.e., cognitive impairment, fatigue, olfactory dysfunction)^17^. PASC is associated with a nearly 4-fold greater risk for mild cognitive impairment ^18^ and epidemiological data have identified a link between COVID-19 and new-onset diagnoses of Alzheimer’s disease and other dementias^19^.

Viral persistence, dysregulated immune responses (e.g., prolonged inflammation, aberrant immune cell activation, and virus-induced autoimmune responses), and tissue reparative defects are the leading hypothesized mechanisms of long-term organ-specific tissue damage during PASC^1,20,21^. Evidence of viral persistence in several settings has been documented^22,23^, but long-term antiviral treatment does not effectively eliminate or improve PASC symptoms in humans^24^. Dysfunctional immune responses, exhaustion, and sustained inflammation long after clearance of infectious virus has been a characteristic in patients and mouse models^25^ with circulating proinflammatory myeloid populations, concentrations of inflammatory biomarkers and cytokines, and dysfunctional T cell subsets consistently reported in clinical studies^26–28^; circulating immune cells, however, represent a small subset of the immune system compared to tissue-resident immune cells that are shaped by local environments. With heterogeneity of symptoms, differential baseline health characteristics in people with PASC, and varying states of immunity, pinpointing the etiologies of PASC has been difficult^25^. This is where animal models are valuable because variables, e.g., baseline health, age, sex, and exposure, can be controlled and genetic mechanisms of PASC can be explored.

To elucidate the biological mechanisms underlying sex differences in PASC, we developed a mouse model and evaluated long-term viral, inflammatory, cognitive and behavioral outcomes. Our study design and analyses involved comparing middle age SARS-CoV-2 infected males and females over time, as well as comparing infected and mock-infected mice within a sex to identify sex differences as well as sex-specific effects of infection. After viral clearance and recovery from acute infection, females exhibit deficits in spatial working and recognition memory, neuroinflammation, and sustained dysregulated pulmonary and systemic immune responses. Using mutant mouse models to dissect the biological factors contributing to differences in disease, we observed that the presence of two X-chromosomes and overexpression of X-linked immune-related genes, including *Tlr7*, drive female-biased PASC neurocognitive phenotypes. Therapeutic inhibition of TLR7 signaling after recovery from acute infection prevents the development of neuroinflammation and neurocognitive PASC in females and provides preclinical evidence for a novel treatment.

## Results

### Middle-aged males suffer worse acute SARS-CoV-2 outcomes than females

To establish a mouse model of sex differences in SARS-CoV-2 pathogenesis, male and female C57BL/6 (inbred) and CD-1 (outbred) mice were intranasally infected with the mu variant (B.1.621) of SARS-CoV-2, with a naturally occurring N501Y spike (S) mutation that binds mouse ACE2^29^ to cause virus replication in the respiratory tract and induce pulmonary immune responses. Mice were infected at 10, 30, or 80 weeks of age, corresponding with young (∼21 human years), middle age (∼37 human years), and old age (∼63 human years) adult populations^30^, and were monitored for acute morbidity for 7 days post-infection (dpi). Among both strains of mice, age-related acute disease severity occurred, with morbidity only apparent among middle and old age mice (**Extended Data Fig. 1A-F**), consistent with previous reports using other SARS-CoV-2 variants (e.g., mouse-adapted SARS-CoV-2 and B.1.351)^31,32^. The greatest sex disparity was observed among 30-week old mice, with males experiencing greater morbidity than females (**Extended Data Fig. 1C, F**) supporting epidemiological data^33^.

To explore sex differences in SARS-CoV-2 disease outcomes, 30-week old male and female C57BL/6 mice were infected (B.1.621) and monitored through 84 dpi (**Fig. 1A**). Males exhibited prolonged acute morbidity, both in terms of mass loss and clinical signs of disease, as compared with females (**Fig. 1B-D**). After infection, females began recovering from acute disease within 4 dpi and males still showed signs of sickness without returning to baseline health until 21 dpi (**Fig. 1E**). Histopathological analyses in lung tissue sections were conducted and at 4 dpi inflammation and exudative lung lesions within blood vessels and in interstitial spaces around blood vessels and bronchiolar airways were recorded, with males having higher semi-quantitative pathology scores, and signs of congestion, inflammation, alveolar fibrin, and cell death than females. Fibrinous exudate with exfoliated epithelial cells and necrotic cellular debris in airway and alveolar airspaces were present during SARS-CoV-2 infection, with females exhibiting residual low levels of hyperplasia, lymphoid aggregation, and inflammation through 42 dpi (**Fig. 1F-G**). Long-term pulmonary pathology is evident, with males exhibiting more severe tissue damage than females early during infection, while females maintaining low-grade residual pulmonary inflammation.

**Figure 1.**
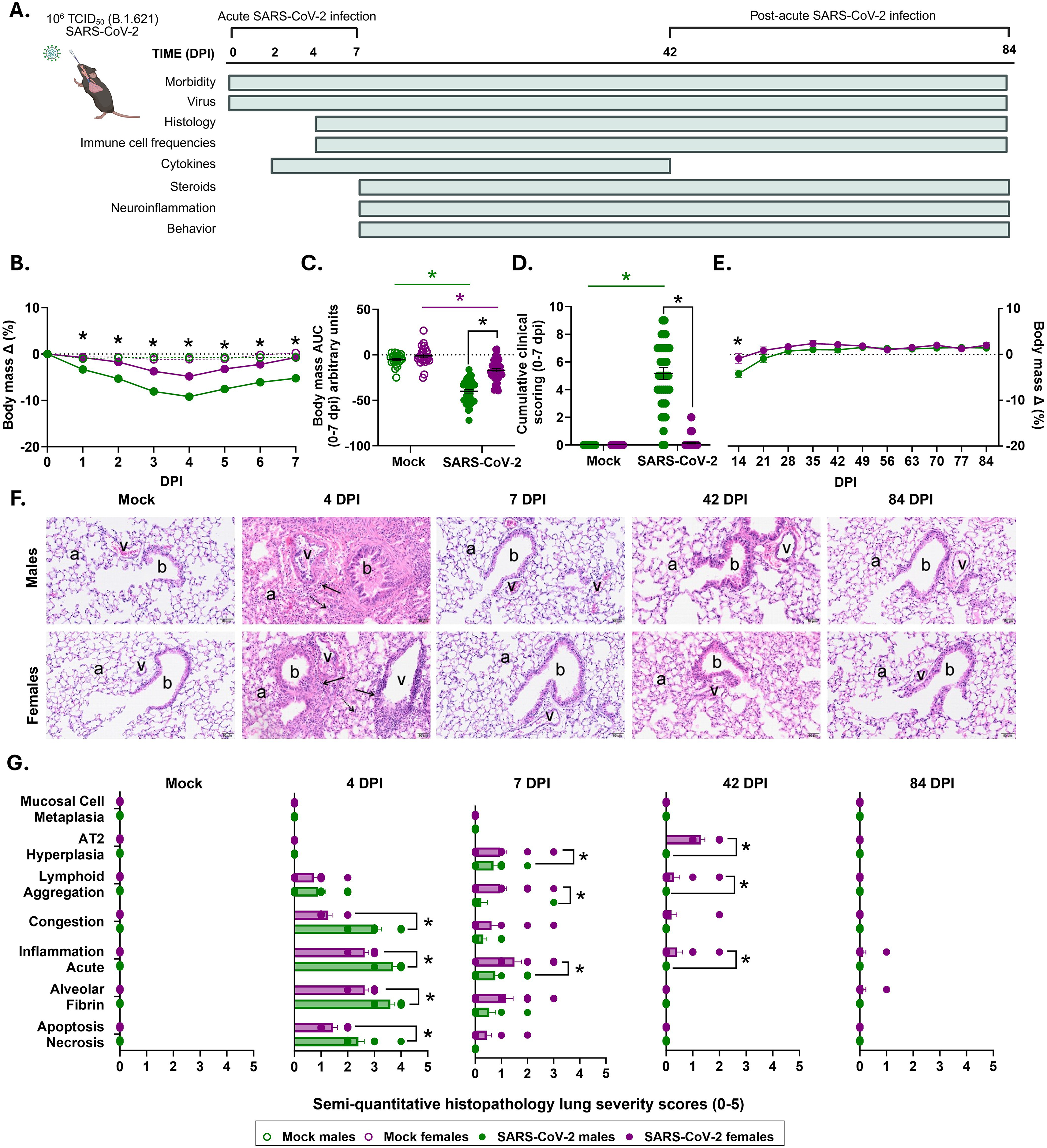
Male mice suffer worse outcomes from acute SARS-CoV-2 than females. Adult male and female C57BL6/J mice were infected with SARS-CoV-2 (B.1.621) or mock-infected at 30 weeks of age, and were monitored through 84 days post infection (dpi) with samples and behavioral indices collected at the specified dpi (**A**). Daily morbidity was recorded and shown as the percent change of body mass from baseline (stippled line) through 7 dpi (**B**). Body mass curves were converted into area under the curve (AUC) values (arbitrary units) for individual mice, where lower AUC values correspond to greater mass loss post infection (**C**). Cumulative clinical disease (**D**), including dyspnea, piloerection, hunched posture, and absence of escape response, were quantified on a scale of 0-4 daily. Body mass continued to be monitored weekly through 84 dpi as a gross indicator of long-term morbidity (**E**). Representative light photomicrographs of tissue sections collected from the left lung lobe stained for hematoxylin and eosin (H&E) collected from mock and SARS-CoV-2 infected males and females through 84 dpi are shown (**F**), scale bar 50 μm, with the following labels: alveolar parenchyma (a), bronchiole (b), blood vessel (v), perivascular inflammatory cells (solid arrow) with mainly lymphocytes and lesser numbers of neutrophils, thickened alveolar septa due to congestion and inflammatory cell infiltrate (stipple arrow). Semi-quantitative pulmonary histopathological assessments were scored as a percentage of total affected tissue (0 = 0%, 1 = 0-5%, 2 = 6-10%, 3 = 11-50%, 4 = 51-75%, 5 = 76+%) through 84 dpi (**G**). Data are represented as mean ± standard error of the mean (SEM), or median ± SEM **(C)** from two to three independent replications (morbidity; n =26-41/group, histology n =6-16/group). Statistical significance was determined by a two-way ANOVAs (repeated measures and ordinary ANOVA) followed by Tukey’s multiple-comparison test or the Kruskal-Wallis test. Asterisks (*) represent significant differences (p<0.05) between groups, with black solid lines reflecting sex differences based on a significant interaction followed by multiple comparisons tests between males and females, and colored solid lines representing differences within sex comparing mock and infected mice (green=males, purple=females). Open circle symbols in the figure represent mock-infected mice and closed circle symbols with filled bars represent B.1.621 infected mice, with colors reflecting sex (green=males, purple=females).

### SARS-CoV-2 infectious virus and viral RNA in diverse tissues do not differ by sex

Infectious virus titers, viral nucleocapsid genomic (gRNA), and subgenomic (sgRNA) RNA were measured in respiratory (**Fig. 2A-I**) and extrapulmonary (**Extended Data Fig. 2A-K**) tissues. Consistent with studies in hamsters^34^, sex differences were not observed in respiratory titers of infectious virus, with complete clearance by 7 dpi (**Fig. 2A-C**). There was no detectable infectious virus in extrapulmonary tissues including in the brain, olfactory bub, and small intestines (**Extended Data Fig. 2A-C**), consistent with previous studies^35^. While sex differences in viral gRNA and sgRNA copies in respiratory tissues were not apparent, low level copies of gRNA and sgRNA persisted through 84 dpi, remaining significantly above mock-infected mice that were used to define assay specificity and background levels (**Fig. 2D-I; Extended Data Fig. 2D-K**). Neither viral gRNA nor sgRNA levels were significantly increased in extrapulmonary tissues, including white adipose tissue, compared to mock-infected counterparts (**Extended Data Fig. 2D-K**). These data suggest that the sexes do not differ in their ability to control viral replication or clear SARS-CoV-2 virus from the respiratory tract, and that low levels of viral RNA persist in respiratory, but not extrapulmonary, tissues.

**Figure 2.**
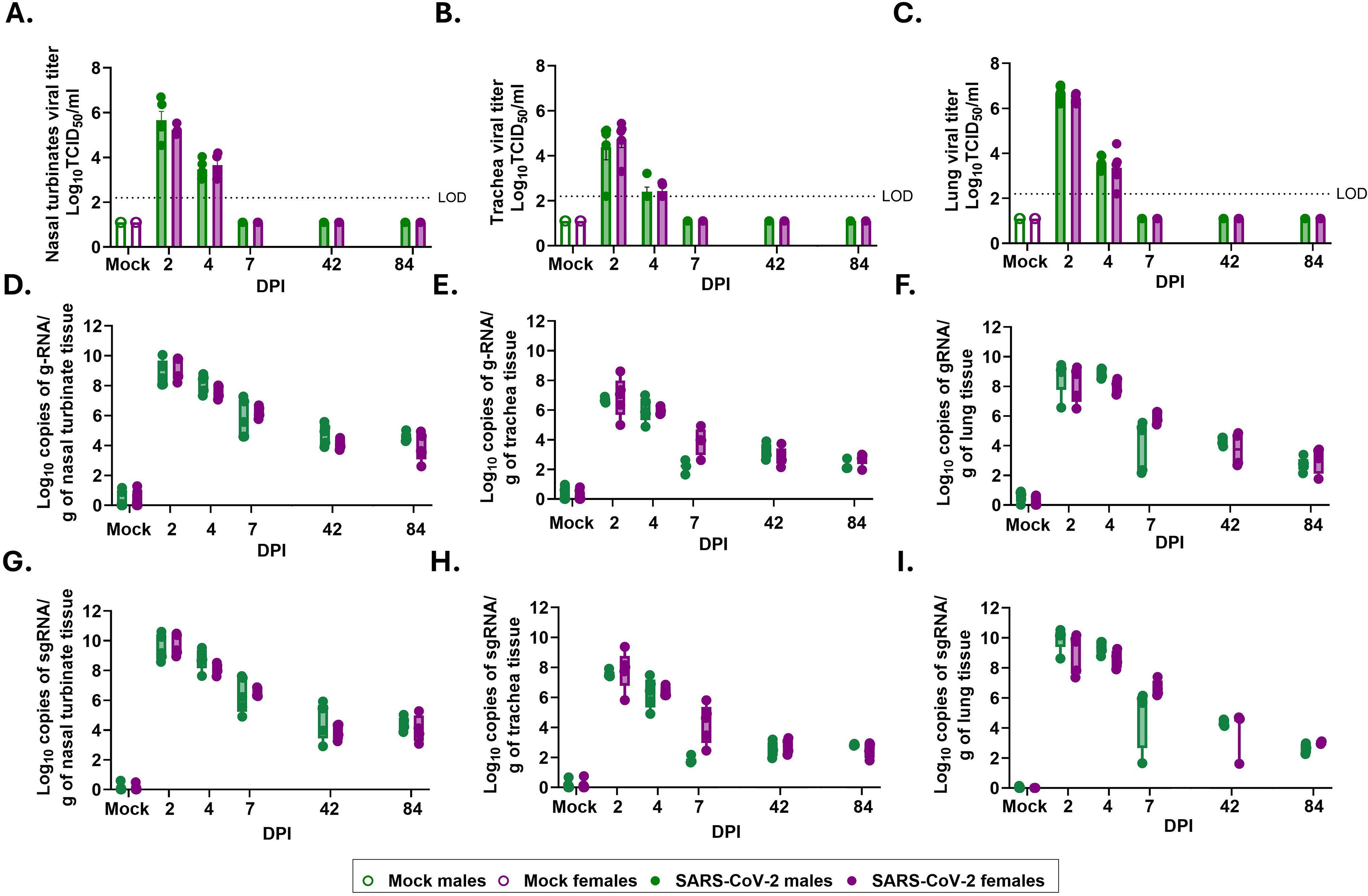
Kinetics of SARS-CoV-2 infectious virus, subgenomic (sg) RNA, genomic (g) RNA in respiratory tissues do not differ by sex. Subsets of SARS-CoV-2 (B.1.621) or mock-infected male and female C57BL6/J mice were euthanized at experimental timepoints, and infectious virus titers as measured by TCID_50_ (**A-C**), subgenomic (sg) viral RNA (**D-F**), and genomic (g) viral RNA (**G-I**) as measured by double droplet PCR were measured respiratory tissues (in nasal turbinates, trachea, and lungs) at 2, 4, 7, 42, or 84 days post-infection (dpi). Data are represented as mean ± SEM from one to two independent replications (virus titers; n =4-5/group; viral RNA n=4-9/group). Statistical significance was determined by a two-way ANOVA followed by Tukey’s multiple-comparison test or Kruskal-Wallis test, followed by the Mann-Whitney test for multiple comparisons. The dashed line indicates the limit of detection (LOD). Open circles/bars represent mock-infected mice and closed circles/bars represent B.1.621 infected mice, with colors reflecting sex (green=males, purple=females).

### Neurocognitive impairments and neuroinflammatory responses are more pronounced in females than males following SARS-CoV-2 infection

To determine whether SARS-CoV-2 infection resulted in long-term cognitive or behavioral changes, we established a battery of behavioral tests (e.g., learning and memory, locomotion, anxiety-like, or depressive-like behaviors) in the animal biosafety level 3 or 2 settings depending on whether experiments were conducted prior to or after 2025 (**Supplemental Materials, Fig. 1**). Our focus was on learning and memory tasks because memory loss is a sex differential characteristic of PASC patients^2,3,5,10,36^. To evaluate distinct domains of cognitive function and memory in mice, we examined spatial navigation, working memory, and conducted a non-spatial recognition task utilizing visual recall.

Spatial working memory was assessed using an spontaneous alternation test that evaluates arm entry visits using video tracking software in an adapted T-maze^37,38^ (**Fig. 3A-C**). This task used rodents’ natural tendency to explore novel, unvisited environments (i.e., arms) in a 3-way maze, using spatial memory recall. A spontaneous alternation was defined as consecutive entries into three distinct arms, and the total percentage of correct spontaneous alternations out of the total number of arm entries was recorded with greater percentages of spontaneous alternations reflecting intact spatial working memory and hippocampal function. As early as 7 dpi (**Extended Data Fig. 3A**) and through 42 and 84 dpi (**Fig. 3B**), infected females had lower percentages of correct spontaneous alternations than either infected males or mock-infected females, with lower percentages indicating cognitive impairment^37^. For behavioral test validation, the number of arm entries was measured to confirm that the reduced percentage of correct alternations was representative of a spatial working memory impairment, and not due to differences in the number of opportunities to complete a three-arm consecutive alternation task. The number of arm entries (**Fig. 3C** and **Extended Data Fig. 3B**) and the total distance travelled in the T-maze (**Extended Data Fig. 3C)** were not affected by either sex or infection.

**Figure 3.**
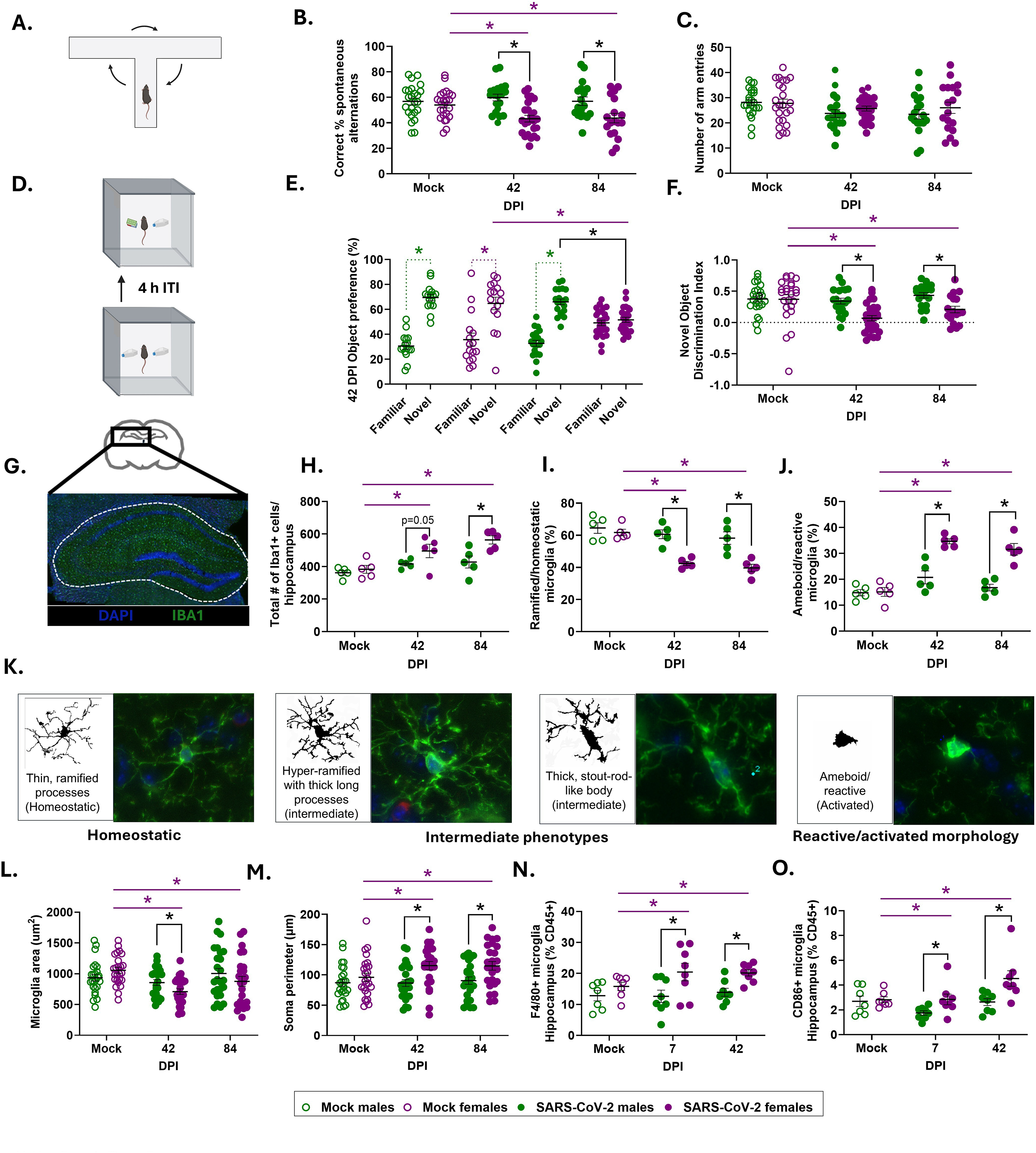
Neurocognitive impairments and neuroinflammatory responses are more pronounced in females than males following SARS-CoV-2 infection. Cognitive function was assessed in SARS-CoV-2 (B.1.621) infected or mock-infected male and female mice at 42 or 84 days post infection (dpi), using a spontaneous alternation task in a T-maze (**A**)as a measure of short-term spatial working memory. In the T-maze task, the percent correct spontaneous alternation was assessed at 42 and 84 dpi (**B**). The percentage of correct spontaneous alternations was calculated based the series of 3 consecutive novel arm entries during testing intervals. Percent alternation was calculated using the following formula: [(# alternations/maximum possible number of alternations) × 100]. Maximum possible alterations were the total arm entries − 2. The total number of total arm entries during the testing interval was quantified (**C**). Novel object recognition (NOR) test was used to test recognition memory by assessing the ability of mice to distinguish familiar (previously presented) objects from a novel object after a 4 h inter-trial interval (ITI); illustrative schematic of NOR (**D**). The object recognition index was calculated by comparing the time spent investigating either the familiar or novel object, reported as a percentage of object preference during the first ∼20 (±2) seconds of total exploration time. Object preference was quantified at 42 dpi (**E**) and novel object discrimination indexes were reported comparing 42 and 84 dpi (**F**). At experimental timepoints, mice were euthanized for either collection of fresh tissue for flow cytometry or perfusion with 1X PBS, followed by 4% paraformaldehyde to preserve brain tissue and structure for immunofluorescence staining **(G)**. Numbers of Iba1+ cells (microglia) in the hippocampus were quantified from immunofluorescence images of brain sections in subsets of SARS-CoV-2 and mock-infected male and female mice at 42 and 84 dpi (**H**). Microglial morphological states (ramified, hypertrophic, rod-like, activated/ameboid) were classified from maximal projected 30 μm z-stack images of microglia in the hippocampus (**I,J**), with quantitative assessments of individual segmented cells (**K**). Morphological parameters and segment characteristics of individual cells, including cell area (**L**) and soma perimeter (**M**) were quantified from binarized cell images. Individual microglial segment characteristic analysis were randomly isolated with an n=25/group performed across 4-5 40X maximal projected Z-stack images in the dentate gyrus, with individual points representing one individual microglia (**L,M**). Single cell suspensions were generated from the hippocampus and immune cells were isolated for flow cytometry of microglial and myeloid phenotypes (**N,O**). Cell populations were gated based on unstained and FMO controls. Data acquisition was conducted using an Cytek Northern Lights spectral flow cytometer, and flow cytometry data were evaluated using FlowJo v10. Data are represented as mean ± SEM from three independent experiments (Behavior; n=8-22/group; immunofluorescence n=5/group, flow cytometry n=6-8/group). Statistical significance was determined by a two-way ANOVA followed by Tukey’s multiple-comparison test, linear or mixed effects model. Asterisks (*) represent significant differences (p<0.05) between groups; black solid lines reflect sex differences in the interaction term followed by multiple comparisons tests between males and females, and colored solid lines represent differences within a sex between mock and/or infected mice at different dpi (green=males, purple=females). Colored stippled lines to represent differences within sexes for object preference. Open circles/bars represent mock-infected mice and closed circle/bars represent B.1.621 infected mice, with colors reflecting sex (green=males, purple=females).

We conducted the novel object recognition (NOR) test to evaluate recognition memory in mice that relies on the hippocampus and medial prefrontal cortex for non-spatial object recognition^39,40^. Following habituation, experimental mice were presented with two identical objects during a 10 min familiarization session and four hours later, mice were returned to the testing environment with one of the two objects replaced with a novel object, and the percentage of time spent investigating either object was quantified (**Fig. 3D**). At 42 dpi, mock-infected mice of both sexes and infected males spent proportionally more time investigating the novel object than the familiar object (**Fig. 3E**), demonstrating a preference for novelty that is indicative of intact recognition memory (**Fig. 3E-F**). Infected females showed no object discrimination or preference for the novel object, indicating an impairment in short-term memory recall. Sex differences in novel object preference indices were apparent at 42 and 84 dpi, with lower scores among infected females as compared to either infected males or mock-infected females (**Fig. 3F**). Object discrimination in females was recovered by 84 dpi, but the percentage of time spent investigating the novel object remained lower in infected compared to mock-infected females (**Fig. 3F** and **Extended Data Fig. 3D**). Together, these data illustrate that in this model, hippocampal-dependent memory deficits are more prevalent in females than males.

Inflammation in the central nervous system (CNS) underlies a number of neurological, neurodegenerative, and cognitive disorders^41^, and has been linked in the development of PASC in animal models and patients^42,43^. With little direct evidence of SARS-CoV-2 virus in the brain, other mechanisms, including neuroinflammation, crosstalk in peripheral and central myeloid populations, blood-brain barrier (BBB) disruption, or downstream impaired neurogenesis may be involved. Broad CNS inflammation was not observed based on concentrations of cytokines and chemokines in whole brain homogenates collected 4-42 dpi (**Supplementary Table 2**). To assess the spatial distribution and morphology of myeloid cell populations in the CNS at post-acute timepoints (**Fig. 3G-J, Extended Data Fig. 3E-H**), immunofluorescent imaging of cells stained with Iba1 was used to identify myeloid immune cells and visualize ramifications and morphology^44^. Co-staining with the microglial-specific marker, TMEM119^45^ was used to determine if myeloid cells were resident or infiltrating cells. Iba1+ cells co-localized with TMEM119 in the hippocampus (dentate gyrus (DG) and CA1-3 regions) confirming their identity as microglia in brain parenchyma. There was no evidence of Iba1+TMEM119- cells in the hippocampus, ruling out the presence of infiltrating macrophages (**Extended Data Fig. 3E**). The absence of peripheral immune cell infiltration was further confirmed by a lack of CD3 (T cells) and Ly6G (neutrophils) immunoreactivity (**Extended Data Fig. 3F**).

Infected females had greater numbers of Iba1+ microglia in the hippocampus compared to infected males or mock-infected females through 84 dpi (**Fig. 3H**). We classified microglia morphology in the hippocampus to define their activation state by visually categorizing cells from branched ramified processes (homeostatic microglia), intermediate hyper-ramified or rod-like phenotypes, to rounder ameboid-like (activated) morphology (**Fig. 3I-K, Extended Data Fig. 3G-H**). Infected females had lower frequencies of ramified homeostatic microglia (**Fig. 3I**) with greater corresponding frequencies of reactive (activated) microglia than infected males or mock-infected females at 42 and 84 dpi (**Fig. 3I-J**). There were no sex differences or effects of infection in intermediate microglia phenotypes (**Extended Data Fig. 3G-H**). Individual microglial cell morphological characteristics^46^ were assessed for quantitative measures of cellular morphology by isolating and binarizing individual cells and perform full skeletal, fractal, and sholl image analyses, with individual data points reflecting independent microglia (**Fig. 3K-O, Extended Data Fig. 3I-M**). During microglial activation, cells rapidly undergo changes in morphology by decreasing extensive ramifications that are critical during surveillance and aggregations to the soma. Cell area, branches, and endpoints per microglia, in addition to span ratio and fractional dimension were reduced in infected females, compared with males (**Fig. 3L, Extended Data Fig. 3J-K)**. Soma perimeter and circularity were increased in individual microglia from infected females compared to males (**Fig. 3M**, **Extended Data Fig. 3L)**, and with no changes in lacunarity (**Extended Data Fig. 3M)**; together, reflecting quantitative features that are reflective of microglial activation. Flow cytometric analysis of the hippocampus further characterized myeloid phenotypes in the brain (**Fig. 3N-O; Gating strategy, Extended Data Fig. 4A**). At 7 dpi, infected females had increased frequencies of microglia and activated CD86+ proinflammatory microglia that remained elevated 42 dpi as compared to either mock-infected females or infected males (**Fig. 3N-O**). Infected females exhibit both neurocognitive impairments (i.e., deficits in spatial working memory and recognition memory) and persistent neuroinflammation.

To determine whether prolonged microglia activation induced broad tissue damage, neurodegeneration, or impaired neurogenesis, immunofluorescent staining of neuronal nuclei (NeuN) and doublecortin (DCX) was used on hippocampal sections. No sex differences or effects of infection were observed on either the percentage of mature neurons or the number of immature neurons at 7-84 dpi (**Extended Data Fig. 4B-D**). NeuN to DCX ratios were used to assess the relative proportion of mature to immature neurons within the hippocampus and revealed no effects of sex or infection at 7, 42, or 84 dpi (**Extended Data Fig. 4E**). ZO-1 was used to identify tight-junction stabilization along blood vessels and epithelial cells in the DG as an indirect measure of BBB permeability (**Extended Data Fig. 4F**). There was no impact of either sex or infection on percentage of ZO-1 positive staining in the DG (**Extended Data Fig. 4G**). Despite the observation of reduced neurogenesis in other mouse models of PASC^35^, neither damage nor impaired neurogenesis in the hippocampus likely underlies cognitive phenotypes in middle-aged infected females compared to males.

### Pulmonary and systemic immune responses are greater in females than males during acute and post-acute SARS-CoV-2 infection

Immune dysregulation and persistent immune cell activation of myeloid and T cell subsets are features observed in individuals with PASC^27^. Cytokine and chemokine concentrations were measured 2-42 dpi and were only elevated in the lungs during peak virus replication (2 dpi) (**Table S1**). At 2 dpi, 35/38 analytes were detectable, with almost half (16/35) of the analytes that were increased at 2 dpi exhibiting sex differences. 15/16 analytes had greater concentrations in infected females than males, including TNF-α, IL-6, IL-1β, CCL2, CCL3, and CCL4, which are involved in myeloid activation and recruitment (**Fig. 4A and Table S1**).

**Figure 4.**
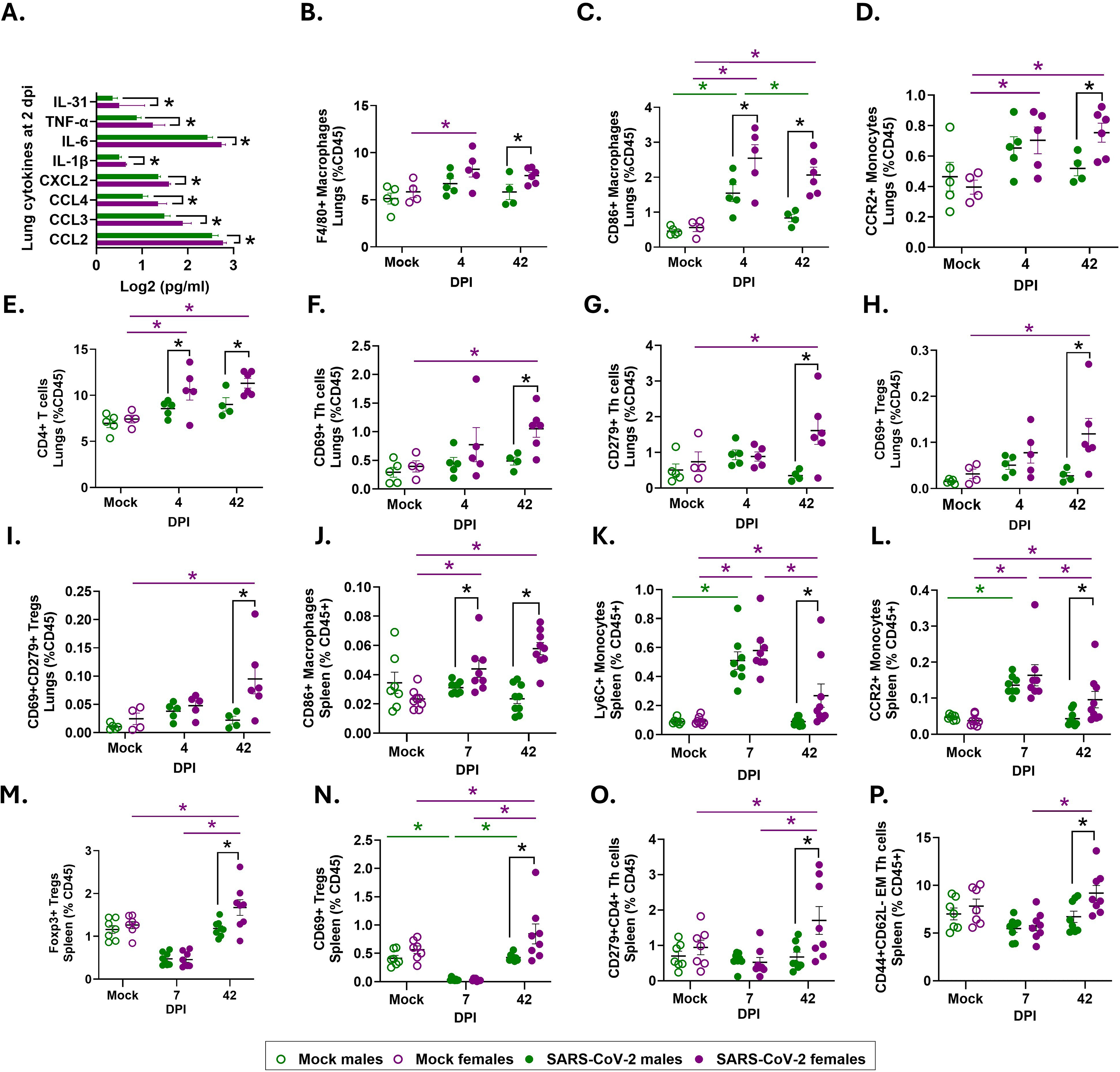
Inflammatory immune responses are greater in females than males during acute and post-acute SARS-CoV-2 infection. Cytokines and chemokines were measured in lung homogenates collected from male and female mice at 2 days post infection (dpi) with SARS-CoV-2 (B.1.621) **(A)**. Single cell suspensions were prepared from the lungs and spleen tissue, myeloid cell subsets and frequencies were analyzed during acute and post-acute infection using spectral flow cytometry. Frequencies of myeloid populations were quantified in the lungs at 4 and 42 dpi. Macrophages **(B)**, CD86+ macrophages **(C)**, CCR2+ monocytes **(D)**, and lymphoid CD4+ T populations, including CD4+ cells **(E)**, CD69+ Th cells **(F)**, CD279+ Th cells **(G),** CD69+ T regulatory (Treg) cells **(H),** and CD69+CD279+ Treg cells **(I),** were measured in mock and SARS-CoV-2 infected males and females, and are reported as a percentage of total CD45+ live cells. Splenic CD86+ macrophages **(J)**, monocytes **(K),** CCR2+ monocytes **(L)**, Tregs **(M)**, CD69+ Tregs **(N)**, CD279+ Th cells **(O),** and CD44+CD62L effector memory Th cells **(P**) were measured at 7 and 42 dpi and reported as a percentage of live CD45+ cells. Data are represented as mean ± SEM from 1-3 independent replications (cytokines; n =4-5/group; flow cytometry n=4-8/group). Statistical significance was determined by a two-way ANOVA followed by Tukey’s multiple-comparison test. Unpaired individual two-tailed parametric t-tests were used to compare individual cytokine/chemokine concentrations at 2 dpi. Asterisks (*) represent significant differences (p<0.05) between groups, with black solid lines reflecting sex differences in the interaction term followed by multiple comparisons tests between males and females, and colored solid lines representing differences within sex comparing mock and infected mice (green=males, purple=females). Open circles/bars represent mock-infected mice and closed circles/bars represent B.1.621 infected mice, with colors reflecting sex (green=males, purple=females).

Spectral flow cytometry was used to characterize myeloid (**Gating strategy, Extended Data Fig. 5A**) and lymphoid (**Gating strategy, Extended Data Fig. 6A-B**) cells in the lungs (site of infection) and spleen. At 4 dpi, when infectious virus was still present in the lungs, frequencies of activated myeloid subsets and neutrophils were elevated in infected compared to mock infected sex-matched mice (**Fig. 4B-D, Extended Data Fig. 5B-D**). Infected females had greater frequencies of F4/80+ macrophages (**Fig. 4B**), pro-inflammatory CD86+ macrophages (**Fig. 4C**), and inflammatory CCR2+ monocytes (**Fig. 4D**) that remained elevated at 42 dpi as compared to either infected males or mock-infected females. Frequencies of all myeloid cell populations returned to baseline in infected males by 42 dpi (**Fig. 4B-D; Extended Data Fig. 5B-D**). Neutrophils were elevated at 4 dpi and returned to baseline by 42 dpi in both sexes (**Extended Data Fig. 5D**). Consistent with other respiratory viruses^47^ ^48,49^, females mounted greater pulmonary inflammatory immune responses to SARS-CoV-2 than males during virus replication, and in many cases these myeloid responses remained elevated long after virus was cleared.

Frequencies of CD4+ T cells were increased in the lungs of infected females compared to males at 4 and 42 dpi (**Fig. 4E**). Many helper T cell subsets, including activated CD69+ Tregs and Th cells, particularly Th cells expressing CD279 (PD-1, exhaustion maker) were increased at 42 dpi in the lungs of infected females compared with either infected males or mock-infected females (**Fig. 4F-I**). Circulating T cell dysregulation and exhaustion is a feature reported in PASC patients^27,50–52^, that also occur in the lungs of female mice.

To determine if sex differences in systemic myeloid and T cell responses also were observed, we measured splenic immune cell populations at 7 dpi, immediately after clearance of infectious virus, and 42 dpi (**Fig. 4J-P; Extended Data Fig. 5E-H, and 7A-S**). In spleens, frequencies of Ly6C+ and CCR2+ monocytes increased 7 dpi in both males and females, but like CD86+ macrophages remained elevated in infected females compared with either infected males or mock-infected females at 42 dpi (**Fig. 4J-L**). Frequencies of splenic F4/80+ and MHCII+ macrophages (**Extended Data Fig. 5E-F**) were greater in infected females than males 7 dpi, but not at 42 dpi. CD206+ macrophages increased at 7 dpi in both sexes and were back to baseline frequencies at 42 dpi, with no sex differences noted (**Extended Data Fig. 5G**). Infected females also had greater proportions of splenic neutrophils and NK cells at 42 dpi than either infected males or mock-infected females (**Extended Data Fig. 5H and 7A-B**).

While infection did not affect total frequences of CD4+ T cells in the spleen (**Extended Data Fig. 7C**), infected females had greater frequencies of activated Tregs and Th cells, including populations of Th cells expressing CD279, CD152 (CTLA4), or CD223 (LAG-3) than either infected males or mock-infected females at 42 dpi (**Fig. 4M-O; Extended data Fig. 7D-H**). There was no notable effect of sex or infection on proportions of CD8+ T cell subsets (**Extended Data Fig. 7I-O**) or B cells (**Extended Data Fig. 7P**). These data demonstrate splenic Th cells from infected females remain activated and exhausted long after SARS-Cov-2 has been cleared.

To determine if memory T cell phenotypes differed between sexes, central memory (CM) and effector memory (EM) subsets were quantified based on CD44 and CD62L expression in splenic CD4+ and CD8+ T cells (**Gating strategy, Extended Data Fig. 6B**). Infected females had lower frequencies of naïve CD4+ cells (**Extended Data Fig. 7Q**), with greater proportions of CM and EM CD4+ T cells than either infected males or mock-infected females at 42 dpi (**Fig. 4P; Extended Data Fig. 7R**). Among CD8+ T cells, there was no impact of infection or sex on frequencies of naïve cells (**Extended Data Fig. 7S**); infected females, however, had greater frequencies of CM and EM CD8+ T cells than infected males at 42 dpi (**Extended Data Fig. 7S-T**). Prolonged activation of myeloid cells, followed by T cell exhaustion and greater T cell memory in infected females compared with males suggests that impaired crosstalk between innate and adaptive immunity in PASC may underlie sex differences in neurocognitive phenotypes of disease.

### ChrX dosage mediates sex differences in acute and post-acute SARS-CoV-2 outcomes

Phenotypic sex differences are mediated by either the direct genetic, indirect gonadal steroid differences caused by sex chromosome complement (i.e., XX vs. XY)^53^ or by an interaction among genes and hormones. Gonadal steroids, including testosterone and estradiol, were measured in serum during acute and post-acute infection. Infection did not alter gonadal steroid concentrations in either sex. Males maintained greater concentrations of testosterone than females (**Fig. 5A**), and estradiol concentrations remained consistent among females, with levels below the limit of detection in males. **(Fig. 5B**).

**Figure 5.**
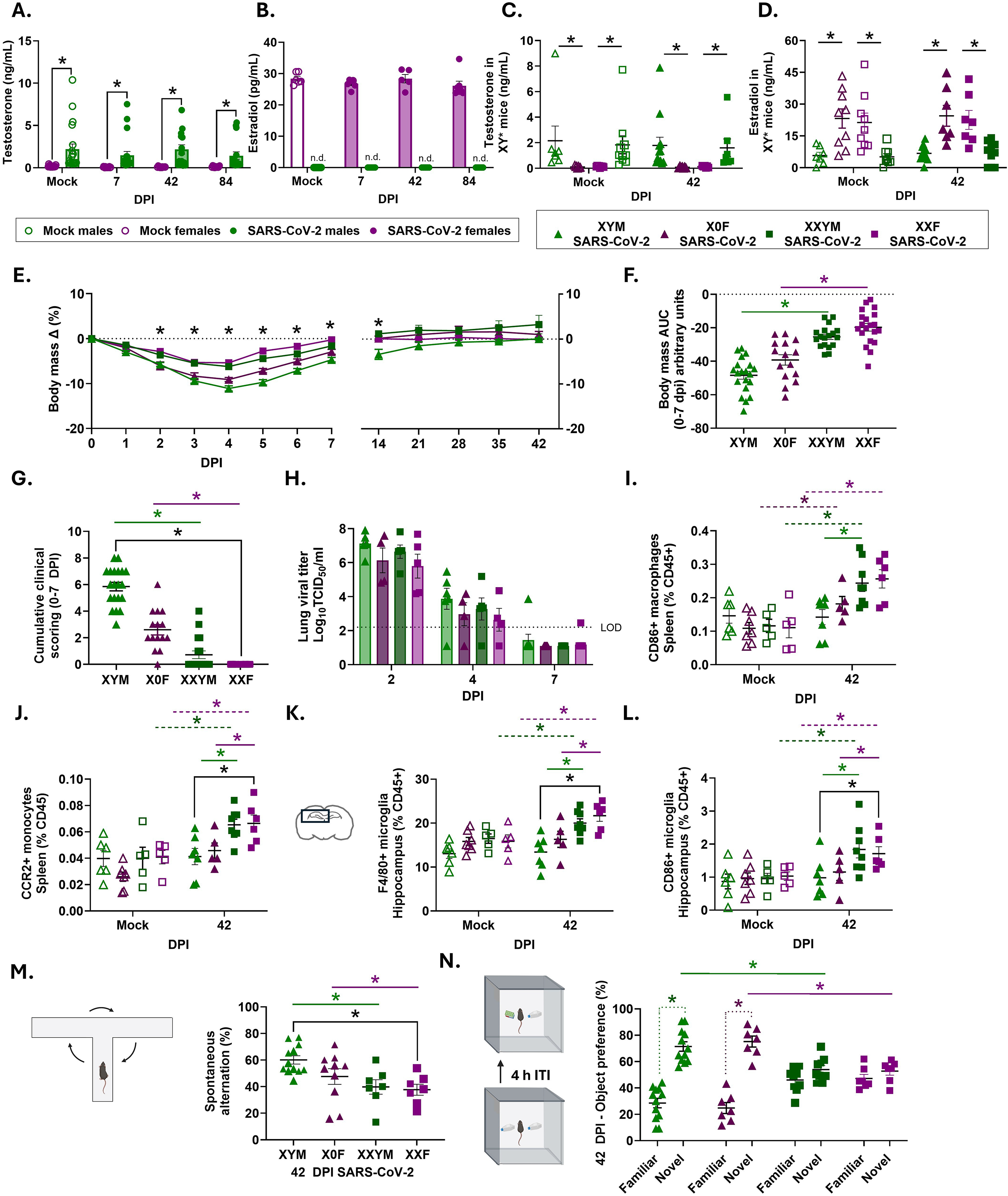
ChrX dosage more than steroids mediate sex differences in acute and post-acute SARS-CoV-2 outcomes. Concentrations of steroids, including testosterone (**A,C**), and estradiol (**B,D**), were measured in serum from infected, and mock-infected experimental wildtype (WT) mice at 7, 42, and 84 dpi, and in XY* mice at 42, or 84 dpi. XY* mice, including gonadal females (XX or X0 chromosome complement) and gonadal males (XY or XXY chromosome complement), were infected with SARS-CoV-2 (B.1.621) or mock-infected at 30-weeks of age and monitored daily for body mass loss (**E-F**) and cumulative clinical disease scores (**G**) through 7 days post infection (dpi) and then weekly until 42 dpi. Infectious virus titers were measured by TCID_50_ in lung tissue through 7 dpi (**H**). The dashed line indicates the limit of detection (LOD). Single cell suspensions were generated from spleens of infected or mock-infected XY* mice and analyzed by flow cytometry at 42 dpi to quantify frequencies of inflammatory CD86+F4/80+ (**I**) and CCR2+Ly6C+ monocytes (**J**). Single cell suspensions were generated from the hippocampus of infected or mock-infected transgenic XY* mice and analyzed by spectral flow cytometry at 42 dpi. Frequencies of F4/80+ macrophages (**K**) CD86+ macrophages and (**L**) microglia reported as a percentage of live CD45+ cells. Cognitive function and spatial working memory were assessed at 42 dpi by measuring spontaneous alternations in a T maze (**M**) and novel object preference in the novel object recognition task (**N**). Data are represented as mean ± SEM, or median ± SEM from two independent replications in wildtype mice (**A-B;** n=5-20/group), and three to six independent replications in XY* mice (morbidity n=7-18/group, infectious virus titers n=4-7/group, flow cytometry n=5-8/group, and behavior n=7-12/group). Statistical significance was determined by ordinary one-way ANOVA (**F**), Kruskal Wallis test (**G**), or a two-way ANOVA (ordinary and repeated measures) followed by Tukey’s multiple-comparison test. Asterisks (*) represent significant differences (p<0.05) between groups; black solid lines reflect sex differences in the interaction term followed by multiple comparisons tests between males and females; colored solid lines represent differences within a gonadal sex (1 vs. 2 chromosomes) **(F-G)**, colored stippled lines representing differences within sex comparing mock and infected mice **(I-L**), and colored dotted lines represent differences within genotype **(N)** (green= males; purple= females). Open circle symbols in the figure represent individual mock-infected WT mice and closed circle symbols with filled bars represent B.1.621 infected mice (**A-B**). Individual data points represent separate experimental mice with triangle symbols representing mice with one X chromosome, square symbols representing mice with two X chromosomes, and color reflecting gonadal sex (green = males, purples = females). Open symbols represent mock-infected mice and closed symbols represent B.1.621 infected mice (**D-N**).

To examine the role of sex chromosomes versus gonadal sex in mediating disease, we used the XY* mice^54,55^, in which XY* males have a modified mutated pseudoautosomal region on the Y chromosome (ChrY; Y*), resulting in abnormal recombination with the X chromosome (ChrX) to generate offspring that are gonadal females that are XX (XXF) or XO (XY^*X^; denoted XOF), or gonadal males that are XY (XYM) or XXY (XX^*Y^; XXYM) (**Extended Data Fig. 8A**). This model was used to determine if sex differences in SARS-CoV-2 pathogenesis were caused by differing X and Y chromosome dosages (i.e., 1 vs 2 ChrX and 0 vs 1 ChrY) in the same gonadal environment, or if differences were driven by gonadal sex. Gonadal steroids were measured and mice with testes (i.e., XYM and XXYM) had greater concentrations of testosterone than mice with ovaries (i.e., XXF and XOF) (**Fig. 5C**), whereas estradiol concentrations were greater in mice with ovaries than in mice with testes (**Fig. 5D**), indicating there were no differences in circulating hormone concentrations between gonadal males or females independent of chromosome number. There was no effect of infection on gonadal steroid concentrations in XY* mice (**Fig. 5C-D**).

XY* mice were mock-infected (**Extended Data Fig. 7B-C**) or infected with SARS-CoV-2 (B.1.621) and monitored for morbidity through 42 dpi (**Fig. 5E**). At 7 dpi, mice with two ChrX (XXF and XXYM) experienced less morbidity and acute clinical disease than animals with a single ChrX (XYM and XOF), regardless of the presence of ChrY or gonadal steroid concentrations (**Fig. 5F-G**). Sex chromosome complement had no effect on the kinetics of virus replication in respiratory tissues, with similar peak virus titers and clearance by 7dpi in all XY* mice (**Fig. 5H, Extended Data Fig. 7D-E**). Infected mice with two ChrX had greater splenic frequencies of macrophages (**Extended Data Fig. 7F**), CD86+ macrophages (**Fig. 5I**), MHC II+ macrophages (**Extended Data Fig. 7G**), and CCR2+ monocytes (**Fig. 5J**) than mice with a single ChrX, regardless of the presence or absence of ChrY at 42 dpi. Infected XXF and XXYM mice also had greater proportions of F4/80+CD11b+ microglia and activated CD86+ microglia in the hippocampus than infected XYM or XOF at 42 dpi (**Fig. 5K-L**), indicating that prolonged activation of myeloid populations was mediated by ChrX dosage.

With a focus on neurocognition and memory at 42 dpi, infected mice with two ChrX, regardless of ChrY (XXF and XXYM), displayed greater spatial working memory impairments as compared with mice that had a single ChrX (XYM, XOF; **Fig. 5M**). There was no effect of sex chromosome complement on working memory in mock-infected mice (**Extended Data Fig. 7H-I**). Infected mice with two ChrX also failed to display NOR, whereas mice with one ChrX displayed intact NOR (**Fig. 5N**) like that of mock-infected mice (**Extended Data Fig. 7I**). To further evaluate mediators of cognitive impairment and test whether circulating sex steroids could account for ChrX dosage effects, separate multiple linear regressions were conducted in XY* infected gonadal males (XYM, XXYM; testosterone model) and infected gonadal females (XXF, X0F; estradiol model) to determine if spatial working memory scores were associated steroid concentrations (testosterone ng/mL or estradiol pg/mL), ChrX dosage (1 vs 2), or the interaction of steroids and ChrX complement were analyzed. Neither testosterone (β = 0.43, 95% CI -3.266 to 4.124; p = 0.81) nor estradiol (β = 0.10, 95% CI -0.6430 to 0.8427; p = 0. 0.77) alone were associated with cognitive outcomes (**Extended Data Fig. 8K-L**). There were no statistical interactions between testosterone and ChrX dosage (β = 1.192, 95% CI - -5.815 to 8.199; p = 0.72) or estradiol and ChrX dosage (β = -0.1986, 95% CI -1.303 to 0.9055; p = 0.70). ChrX dosage was associated with spatial memory scores in both models (Testosterone β = -21.10, 95% CI -33.93 to -8.274, p = 0.003; and estradiol β = -20.45, 95% CI -33.11 to -7.791, p = 0.005). In summary, infected mice with two ChrX scored lower than mice with one ChrX, illustrating that sex differences in PASC memory deficits were primarily mediated by ChrX dosage, rather than gonadal steroids alone.

### SARS-CoV-2 infection increases expression of ChrX-linked genes

ChrX is enriched for immune response genes. XX females undergo X chromosome inactivation (XCI) to silence one of two ChrX to maintain similar levels of X-linked protein expression between sexes. A subset of X-linked genes (e.g., *Kdm6a*) escape XCI and have higher expression in XX than XY cells^56^. In humans, females that develop PASC have increased expression of the long non-coding RNA, *Xist*, in several immune cells as compared to females who recover from COVID-19^28,57^. *Xist* acts as a ChrX dosage mediator, with any up or downregulation in expression resulting in downstream differences in ChrX gene expression^58^. Using pulmonary macrophages and lung tissue from infected WT and XY* mice, the expression of several ChrX genes was analyzed during acute and post-acute timepoints after SARS-CoV-2 infection. At 2 dpi in WT mice, *Xist* expression was increased in pulmonary macrophages from infected compared to mock-infected WT females and was not expressed in WT males **(Fig. 6A)**. SARS-CoV-2-induced expression of *Xist* remained elevated in pulmonary tissue from WT females through 42 dpi as compared with mock infected females (**Fig. 6B**). The greater infection-induced *Xist* expression in the lungs was limited to XY* mice that had two ChrX in the absence of ChrY (XXF only). The presence of ChrY and/or testes (XXYM) mitigated infection-induced *Xist* expression at 42 dpi (**Fig. 6C**). This phenotype could be due to an interaction between ChrX dosage and gonadal steroids or indirect effect of elevated testosterone on the expression of ChrX genes.

**Figure 6.**
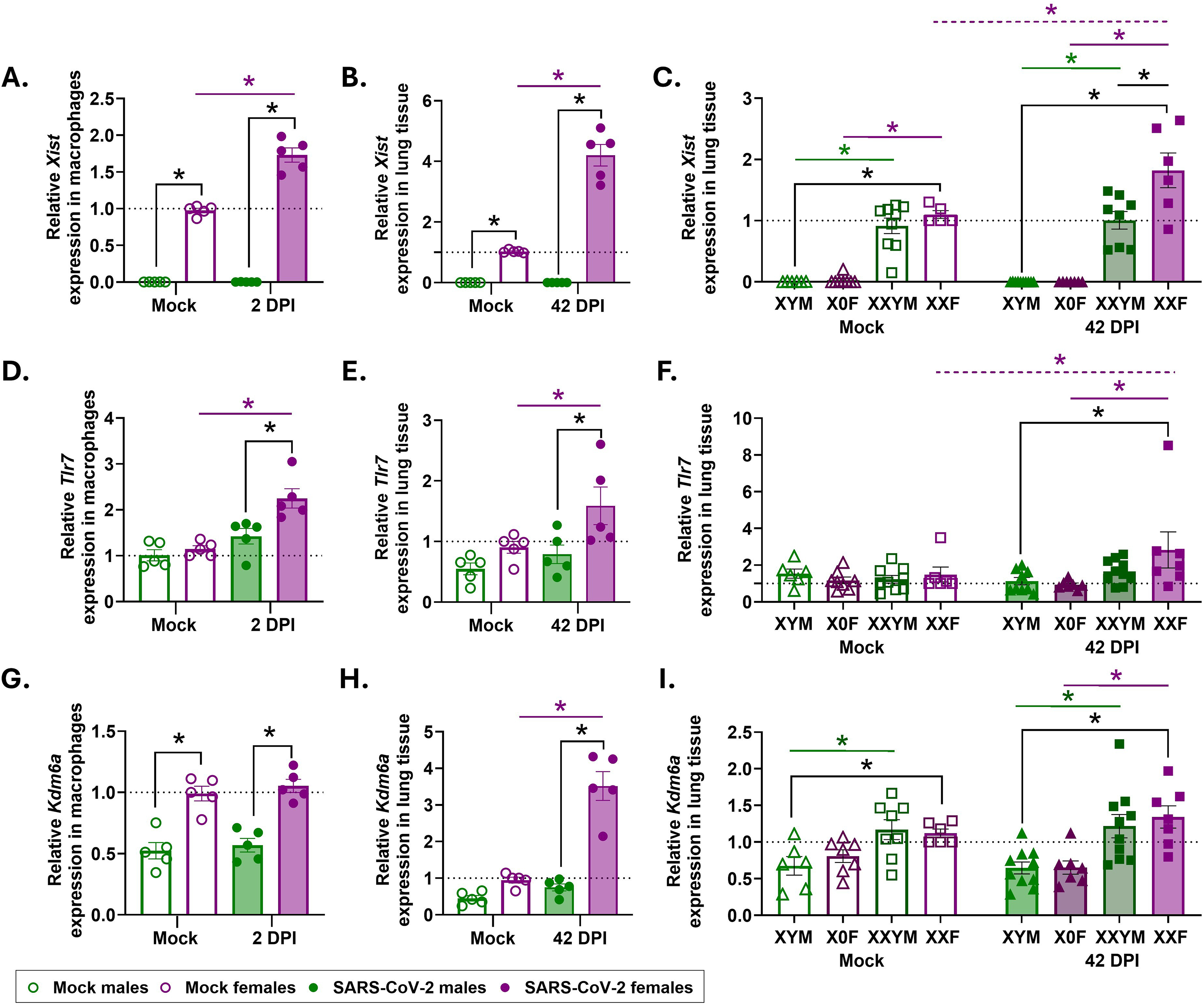
SARS-Cov-2 infection increases the expression of X-linked genes in cells and tissues from XX female mice at acute and post-acute timepoints. Male and female wildtype mice or XY* mice were infected with SARS-CoV-2 (B.1.621) or mock-infected, and euthanized at 2 or, 42 days post infection (dpi) for tissue collection. RNA was isolated from pulmonary F/80+ macrophages at 2 dpi in wildtype mice, from lung homogenates at 42 dpi from wildtype mice, or from lung homogenates at 42 dpi from transgenic XY* for qR-PCR analyses of *Xist* (**A-C**), *Tlr7* (**D-F**), and *Kdm6a* (**G-I**). Relative expression was normalized to expression in uninfected XX females for each gene, with β-actin as a housekeeping control. Data are represented as mean ± SEM from two independent replications in wildtype mice and two to three independent replications in XY* mice (macrophages n=5/group, lungs n=4-11group). Statistical significance was determined by a two-way ANOVA followed by Tukey’s multiple-comparison test. Asterisks (*) represent significant differences (p<0.05) between groups; black solid lines reflect sex differences in the interaction with multiple comparisons tests used to determine significance between males and females; colored solid lines represent differences within sex comparing mock and infected mice (green=males, purple=females). Open circle symbols in the figure represent individual mock-infected WT mice and closed circle symbols with filled bars represent B.1.621 infected mice (**A-B, D-E, G-H**). Triangle symbols represent mice with one X chromosome, and square symbols represent mice with two X chromosomes with color reflecting gonadal sex (green = males, purples = females). Open symbols represent mock-infected mice and closed symbols represent B.1.621 infected mice (**C,F,I**).

**Figure 7.**
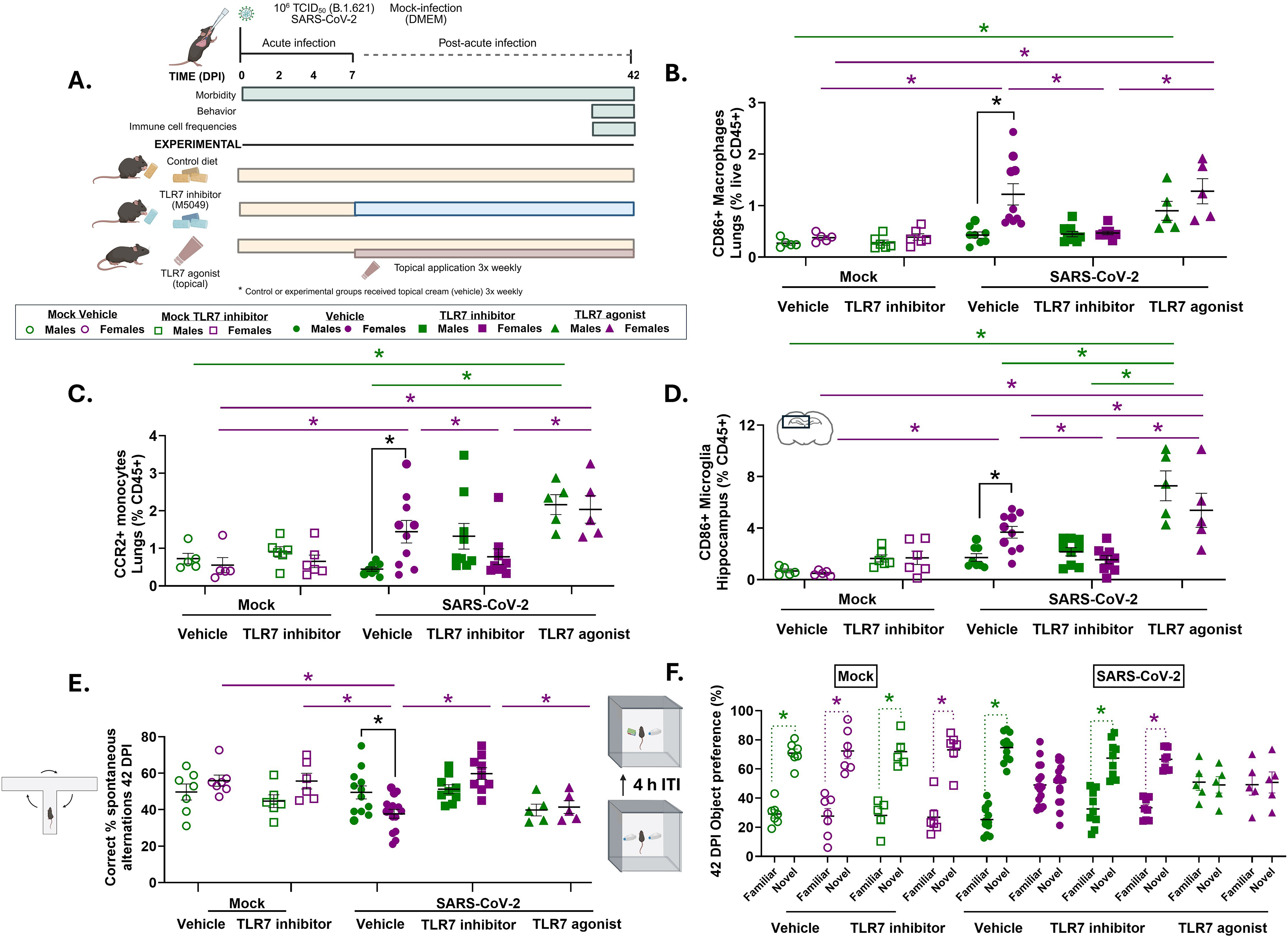
Prolonged TLR7 signaling causes female-biased inflammation and memory loss following SARS-CoV-2 infection. Adult male and female C57BL6/J mice were infected with SARS-CoV-2 (B.1.621) or mock-infected at 30 weeks of age. After 7 days post infection (dpi), male and female mice received either a TLR7 inhibitor (M5049) at 40 mg/kg incorporated into standard rodent chow (SARS-CoV-2 TLR7 inhibitor), or were maintained on batch-matched control diets (SARS-CoV-2 vehicle) for 35 days post-infection with behavior and experimental collection at 42 dpi. For an experimental control at 7 dpi, a subset of mock-infected male and female mice received TLR7 inhibitors (M5049) at 40 mg/kg incorporated into standard rodent chow (Mock TLR7 inhibitor), and for baseline sex-matched comparisons, mock-infected males and females were maintained on batch-matched control diets for the duration of the study (Mock vehicle). A subset of SARS-CoV-2 infected male and female mice were administered with a topical TLR7 agonist (SARS-CoV-2 TLR7 agonist), after 7 dpi. SARS-CoV-2 TLR7 agonist experimental mice were epicutaneously administered 50 mg of 5 % imiquimod cream, with all other groups administered a vehicle cream control. Cognitive assessments were conducted prior to tissue collection at 42 dpi (Experimental Design; **A**). Single cell suspensions were prepared from the lungs and analyzed by spectral flow cytometry at 42 dpi. The frequency of CD86+ macrophages were reported as a percentage out of total macrophages **(B)**. CCR2+ monocytes **(C)** and CD86+ microglia **(D)** were reported as the percentage of total live CD45+ cells. Cognitive function and spatial working memory were assessed by measuring spontaneous alternations in a T maze (**E**) and novel object preference in the novel object recognition task (**F**). Data are represented as mean ± standard error of the mean (SEM) from three independent replications (morbidity; n =/group). Statistical significance was determined by a two-way ANOVAs followed by Tukey’s multiple-comparison test. Asterisks (*) represent significant differences (p<0.05) between groups; black solid lines reflect sex differences in multiple comparisons tests between males and females following a significant interaction; colored solid lines represent differences within sex comparing mock and infected mice (green=males, purple=females). Open symbols in the figure represent mock-infected mice and closed symbols represent B.1.621 infected mice. Circle symbols represent vehicle experimental mice, square symbols represent TLR7 inhibitor groups, and triangle symbols represent the TLR7 agonist group, with colors reflecting sex (green=males, purple=females).

*Xist* RNA can also act as a female-specific TLR7 ligand^59^ that may contribute to prolonged immune system activation, as seen in autoimmune diseases^60^. Consistent with *Xist* expression, SARS-CoV-2 infection increased expression of *Tlr7* in pulmonary macrophages at 2 dpi (**Fig. 6D**) and in pulmonary tissue from XX WT females at 42 dpi (**Fig. 6E**). In XY* mice, greater expression of *Tlr7* in pulmonary tissue at 42 dpi was mediated by the presence of two ChrX in the absence of ChrY/testes (XXF) because the addition of ChrY/testes (XXYM) lowered *Tlr7* expression to levels seen in animals with a single ChrX in either the presence (XYM) or absence (XOF) of ChrY (**Fig. 6F**). Dysregulated *Xist* expression could alter XCI and X-linked gene expression, including *Kdm6a* which is a histone demethylase that mediates activation and repression of histone modifications, is known to escape XCI, and is expressed at higher levels in XX compared to XY females^61^. As expected, relative expression of *Kdm6a* was elevated in XX compared to XY macrophages at 2 dpi, but expression levels were not changed by infection (**Fig. 6G**). SARS-CoV-2 infection did, however, increase *Kdm6a* expression in lungs from WT female mice compared with WT male mice, but not in XY* mice at 42 dpi (**Fig. 6H**). The infection-induced increase in *Kdm6a* expression was not replicated in XY* mice, which may be due to effects of ChrY (**Fig. 6I**). These data illustrate that SARS-CoV-2 infection causes prolonged induction of *Xist* and downstream *Tlr7* in XX mice, which is associated with greater inflammation and memory loss associated with PASC.

### Prolonged TLR7 activation causes female-biased inflammation and memory loss associated with PASC

To determine whether increased activation of TLR7 was a cause of neurocognitive sequelae (i.e., spatial working memory impairment and recognition memory deficits) in females, we pharmacologically manipulated TLR7 signaling after recovery from acute infection in wild-type mice (**Fig. 8A**). Mice were infected with SARS-CoV-2 or mock infected (**Extended Data Fig. 9A-B**). Beginning at 7 dpi, mice were administered a selective TLR7/8 inhibitor, Enpatoran (M5049) or a TLR7 agonist, imiquimod (R837) through 42 dpi. Manipulation of TLR7 signaling after 7 dpi did not alter body mass in either male or female mice highlighting that the doses selected did not make the animals sick (**Extended Data Fig. 9C**). To confirm the treatments altered TLR7 signaling, *Irf7* was measured in lung tissue (**Extended Data Fig. 9D**) and was shown to be downregulated by TLR7 inhibition and upregulated by TLR7 stimulation in both infected males and females.

In the lungs, receipt of the TLR7 inhibitor reduced and receipt of the TLR7 agonist increased frequencies of F4/80+ macrophages (**Extended Data Fig. 9E**), CD86+ macrophages (**Fig. 8B, Extended Data Fig. 9F**), and CCR2+ monocytes (**Fig. 8C**) in both infected males and females at 42 dpi relative to their infected, vehicle treated counterparts. Treatment with a TLR7 agonist increased pulmonary neutrophils in infected males and females compared to sex-matched counterparts (**Extended Data Fig. 9G**). Frequencies of activated CD86+ microglia in the hippocampus also were reduced by TLR7 inhibition and increased by TLR7 stimulation in both infected males and females at 42 dpi (**Fig. 8D**). The effects of TLR7 manipulation on myeloid cell populations were limited to infected and not mock-infected males and females. Inhibition of TLR7 signaling activity eliminated the prolonged pulmonary and hippocampal inflammation in infected females. The effects of TLR7 inhibition on myeloid cell populations was consistently greater for infected females than males because males inherently have lower TLR7 signaling than females (**Fig. 6E**).

To determine if elevated TLR7 activity caused neurocognitive sequelae PASC in females, both spontaneous alternations in a T-maze (**Fig. 8E and Extended Data Fig. 9H**) and NOR (**Fig. 8F and Extended Data Fig. 9I**) were measured at 42 dpi. Inhibition of TLR7 activity after recovery from acute SARS-CoV-2 disease prevented spatial working and recognition memory deficits in infected females. TLR7 inhibition resulted in greater percentages of correct spontaneous alternations (**Fig. 8E**) and improved novel object preference (**Fig. 8F**) in infected females, to levels that were comparable with infected males and mock-infected females. TLR7 stimulation, reduced percentages of spontaneous alternations and novel object preference in infected male and female mice to levels comparable with infected females treated with vehicle (**Fig. 8E-F, Extended Data Fig. 9H-I**). Together, these data illustrate that prolonged activation of TLR7 signaling underlies female-biased immune dysregulation and neurocognitive PASC.

## Discussion

COVID-19 was identified as the leading factor associated with the widening life expectancy gap between males and females between 2019-2021^62^. It has been more straight forward to quantify the public health consequences of sex differences in acute COVID-19 than to quantify the sex differential effects of PASC given its 200+ recognized symptoms^36^. Because PASC encompasses many disorders, we focused on the mechanisms of sex differences underlying neurocognitive sequalae, because data consistently show that women are more likely than men to present with brain fog, memory loss, and broad cognitive deficits associated with PASC and other post-acute infection syndromes (PAIS)^63^. In addition to PAIS, women also are more likely than males to experience cognitive symptoms associated with dementia and Alzheimer’s^64–67^.

In our mouse model, males developed worse acute disease compared to females after SARS-CoV-2 infection, consistent with reports in humans^68^ and other rodent models^34^, and highlighting that male-biased severity of acute SARS-CoV-2 is conserved across species and virus variants. After mild-to-moderate infection with B.1.621 middle-aged females have greater inflammatory immune responses following infection that are protective against acute disease but also mediate persistent neuroinflammation and neurocognitive sequelae long after infectious virus has been cleared. This model reflects sex differences across acute and post-acute timepoints, allowing for direct comparisons between sexes and changes over time after recovery, to characterize early immunologic changes that initiate and maintain long-term clinical sequelae. We used this model to demonstrate that female-biased development of long-term pulmonary inflammation, neuroinflammation, and neurocognitive deficits in memory associated with PASC is mediated by a double dosage of ChrX. Infection-induced overexpression of X-linked gene, *Tlr7*, and the prolonged activation of TLR7 signaling, particularly in inflammatory myeloid cells in the lungs and hippocampus. Males can experience female-like memory deficits associated with PASC by either having two ChrX or through prolonged TLR7 activation.

Other models of PASC have been developed and include SARS-CoV-2 infection of humanized ACE2 (hACE2) mice^69,70^ or infection of wild-type mice with other viruses that cause chronic disease^71,72^. Infection of hACE2 mice with SARS-CoV-2 causes severe pathology, acute infection of the CNS, and death, which makes long-term studies more complicated. Prior studies using the beta variant (B.1.351) or mouse-adapted strains report that SARS-CoV-2 infection causes neuroinflammation and dysregulated myeloid responses in the brain, in addition to pulmonary complications, thereby establishing a link between respiratory infection and the CNS, despite no presence of virus in the brain^35,73^, consistent with our mouse model. In the absence of respiratory infection, intracerebral inoculation of SARS-CoV-2 spike protein directly into the brains of mice causes neuroinflammation and cognitive dysfunction, similar to the current study, but without analysis of sex differences^74^. Infection of C57BL/6 mice with mouse hepatitis virus causes deficits in spatial working memory combined with gliosis in females but not males^75^. When sexes of mice have been compared and sex differences, particularly in acute disease, have not been observed, it appears to be due to differences in age (young vs. middle age)^32,35^ or use of more virulent viruses^76^.

To investigate the complexities that underlie the female bias in PASC identified in human clinical data, preclinical models where variables can be controlled are essential to isolate direct effects. We demonstrate that ChrX dosage causes sex differences in acute and post-acute SARS-CoV-2 phenotypes. Concentrations of sex hormones and the presences of either testes or ovaries in XY* mice did not associate with differential inflammatory immune responses or cognitive outcomes. Instead, the patterns of acute disease and prolonged inflammation and cognitive dysfunction following infection depended on the ChrX dosage. Having two ChrX, independent of gonadal sex, gonadal steroids, or presence or absence of the Y chromosome was protective against acute disease, but mediated development of post-acute symptoms in XY* mice. We cannot rule out steroidal effects that may contribute to sex chromosome-mediated disease outcomes through changes in hormone receptor sensitivity, post-receptor signaling pathways, feedback loops, or organizational effects^77^. Our data provide novel converging evidence that ChrX dosage is a primary mediator of sex differences in cognitive PASC outcomes.

Prolonged inflammation and neurocognitive symptoms in females were mediated by SARS-CoV-2 increasing expression of the X-linked genes, *Xist* and *Tlr7* in females, consistent with observations in female patients with either PASC^57,78^ or autoimmune diseases^79^. ChrX mediation of female-biased conditions, including autoimmune diseases, is maintained in males with Klinefelter syndrome (XXY) and females with triple X syndrome (XXX), but not in females with Turner syndrome (XO)^80^. Dysregulated *Xist* expression can impair XCI, lead to overexpression of X-linked genes from the inactivated ChrX and precipitate the development of overactive immune responses that cause chronic conditions^81^. Whether the XIST complex serves as a ligand for TLR7 to cause prolonged inflammatory signaling during PASC in females requires investigation.

While TLR7 is essential for early immune responses against SARS-CoV-2, sustained signaling contributes to sex-based differences in pathogenesis. Deficiencies of *Tlr7* (Tlr7^-/-^) or *Irf7* (Irf7^-/-^) increase the severity of acute SARS-CoV-2 infection, resulting in greater pulmonary pathology and viral loads in male and female mice infected with mouse adapted SARS-CoV-2^82,83^, with higher rates of mortality seen only in males^82^. TLR7 is necessary for viral clearance and recovery, with complete mortality observed in mice where type 1 IFN signaling is blocked using IFNAR blocking antibodies^83^. Male COVID patients with loss of function TLR7 variants developed more severe acute COVID-19 than either their female relatives or males without this variant during the pandemic^84^. There is clear evidence that TLR7 signaling is necessary to recognize and respond to SARS-CoV-2 viral RNA. The SARS-CoV-2 nucleocapsid (N) protein potentiates TLR7-mediated proinflammatory immune responses in macrophages in cell culture^85^. Whether females have greater systemic responses to SARS-CoV-2 N protein that potentiate TLR7-mediated activation of myeloid cells requires consideration. We show that prolonged TLR7 signaling after virus has been cleared from the respiratory track contribute to sustained pulmonary and CNS inflammation and causes development of female-biased neurocognitive PASC that can be pharmacologically reversed.

Persistent type I IFN signaling has been reported in PASC cohorts^86–89^ and pDCs from XX females produce greater IFNα following TLR7 stimulation^90^. Enhanced type I IFN production, induction of NF-κB and TLR7-IRF5 downstream signaling cascades, and pro-inflammatory cytokine and chemokine production (e.g., CCL2, IL-6, TNFα, IL-1β) could be involved and require further consideration. Elevated expression of *Xist*, observed in infected females mice and in human female PASC patients^25^ may also contribute to a TLR7-XIST feedback loop that results in chronic inflammation. *Xist* RNA can act as a ligand for TLR7 in XX cells^59^. Altered *Xist* expression increases escape of X-linked genes^91^ that also may sustain inflammation. Perturbed XCI in macrophages could impair negative feedback mechanisms of inflammation and sustain expression of target genes, directly linking altered XCI to chronic inflammation^91^. Other immune-related genes that escape XCI can form immunoregulatory networks that act synergistically with TLR7 in immune cells to enhance cytokine release in autoimmune conditions^92,93^. Epigenetic remodeling from histone demethylases including *Kdm6a* that escapes XCI can remove repressive H3K27me3 sites and alter methylation patterns to keep inflammatory loci accessible following TLR7 activation, which can facilitate *Xist* upregulation ^94^. Together, these proposed mechanisms offer possible explanations for how acute infection could result in long-term sex-biased inflammatory phenotypes. Further investigation into molecular mechanisms are necessary for post-infectious cognitive impairment with shared pathways to understand immune contributions to female-biased chronic conditions. Our data establish ChrX dosage as a determinant of sex differences in PASC and identify TLR7 as a therapeutic target.

TLR7 inhibitors are currently in clinical trials for autoimmune diseases^95,96^ such as systemic lupus erythematosus^97^, and in preclinical models of Alzheimers^67^. Our findings demonstrate that pharmacological inhibition of TLR7 after recovery from acute SARS-CoV-2 prevents post-acute pro-inflammatory myeloid responses in the lungs and CNS in infected females. Because a systemic inhibitor was used, whether these are direct effects on myeloid cells or indirect effects on other cells requires consideration. Defining the precise cell-specific contributions of TLR7 signaling remains an important direction for future studies using conditional knockout models.

Lastly, age is a critical factor in the presentation of sex differences in PASC both in humans and in our animal model. Clinical studies highlight that middle-aged women, regardless of menopause status, are more likely than middle-aged men to develop PASC^98^. Our data further highlight that gonadal steroid signaling is not attributed to sex differences in neuroinflammation and neurocognitive PASC in mice. The double dosage of ChrX and elevated expression of *Tlr7* in immune cells from females is not age-dependent. The DNA methylation patterns, gene expression profiles, and morphology of microglia in the hippocampus, however, are age-dependent. Microglia in the hippocampus of human autopsy samples (no sex disaggregation) undergo an epigenetic transition in middle age from having characteristics of embryonically-derived, brain-resident microglia to having epigenetic features of blood-derived monocytes with increased soma size and expression of immune responses genes characteristic of neuroinflammation^99^. In addition to making the hippocampus more receptive to age-related cognitive decline, it could be that the layering of infection-induced TLR7 activation in myeloid-derived cells, including monocytes and microglia, on top of the middle age-related epigenetic modification in microglia increases the inflammatory properties of microglia more so in females than males, which requires further investigation. Collectively, the double dosage of ChrX and prolonged activation of TLR7 makes the female hippocampus more vulnerable than the male hippocampus to neuroinflammation, which might explain the preponderance of neurocognitive symptoms in females with PASC and even other PAIS and autoimmune diseases.

## Acknowledgments

We thank the animal care veterinarians and staff from the Miller Research Building Animal Facilities at Johns Hopkins for assistance with maintaining animal colonies. We thank Jamie Perry for assistance with collections, Kayla Kimble and Garrett Chen for assistance with virus quantification, Joe Hoffmann for assistance with flow cytometry, Naomi Esrig for assistance with digital droplet PCR, and Ryan Lewandowski for assistance with digital histology scans. We thank Drs. Montserrat C. Anguera at the University of Pennsylvania, and Nuriban Valero-Pacheco, Vivianne Morrison from the Engler-Chiurazzi and Zwezdaryk lab, along with members of the Klein, Davis, Thompson, and Pekosz labs for discussions about these data. Light microscopy images were generated using the instruments and support of the Light Microscopy Core of the Department of Molecular Microbiology and Immunology at the Johns Hopkins Bloomberg School of Public Health. Illustrative diagrams and schematics were created using Biorender.com. Diagrams in figures were created in BioRender. Liu, J. (2026) https://BioRender.com/fwfwu3e.

## Conceptualization

J.A.L., A.L.C., A.P., S.L.K, **Methodology:** J.A.L., H.S.P., J.R.H., C.L.T, A.B., A.P., APA, R.B., E.E-C., K.J.Z., E.A.T., **Investigation:** J.A.L., S.C., X.X., T.Z., M.T., K.S., P.S.C., W.Z., N.B.B., H.H., **Visualization:** J.A.L., J.R.H., M.T., **Funding acquisition:** S.L.K., A.P., A.L.C., A.P.A. **Project administration:** S.L.K., **Supervision:** S.L.K., **Writing – original draft:** J.A.L., S.L.K., **Writing – review & editing:** All authors.

## Online Methods

### Viruses and cells

The mu B.1.621 (hCoV-19/USA/MD-HP06587/2021) SARS-CoV-2 variant originated from a human clinical isolate (hCoV-19/USA/MD HP06587/2021). The mu variant SARS-CoV-2 virus (hCoV-19/USA/MD HP06587/2021; Pango Lineage B.1.621, GISAID accession number EPI_ISL_3101473) was isolated at Johns Hopkins University as described^100,101^, is available from the Biodefense and Emerging Infections Research Resources Repository (BEI) Resources (catalog number NR-56225), and was originally contributed by Dr. Andrew S. Pekosz. Working stocks of the mu-variant of SARS-CoV-2 were generated by infecting Vero-E6-TMPRSS2 (Japanese Collection of Research Bioresources Cell Bank no. JCRB1819) cells at a multiplicity of infection (MOI) of 0.01 tissue culture infectious dose 50 (TCID50) per cell in infection media (DMEM) that is supplemented with 2.5% filter-sterilized FBS, 100 U/mL penicillin and 100 μg/mL streptomycin, 1 mM l-glutamine, and 1-mM sodium pyruvate. 72 hours following infection, supernatant was collected, centrifuged at 400g for 10 minutes, concentrated, and stored in aliquots at –80°C for infection, as previously described^102^.

### Experimental mice

Adult male and female C57BL/6J mice were purchased from Jackson Laboratories, CD-1 mice were purchased from Charles River Laboratories and if needed, were aged in house to 17 months. Aged (17 month) C57BL6/CR mice were obtained from the National Institute on Aging animal facilities. XY* breeding pairs on a C57BL/6J background (MMRRC strain 043694-UCD) were obtained from Arthur Arnold (University of California, Los Angeles, USA) and mouse lines were maintained in house by mating XY* male mice that contain an altered pseudoautosomal region (PAR) on Chromosome Y (denoted Y*, composed of complete PAR, no NPY, and a small subsegment of NPX)^103^ with wildtype females on a C57BL/6J background to produce offspring that were: gonadal females (XX and X0 chromosomes; XXF, X0F) and gonadal males (XY and XXY chromosomes, XYM, XXYM). Genotypes were confirmed at or after weaning by digital polymerase chain reaction (PCR) analysis for the number of TLR7 and RPP30 copies.

Mice were housed five per cage and maintained under standard animal biosafety level 2 (ABSL-2) until mice reached the appropriate age for infection and were transferred into ABSL-3 facilities or were directly received from vendors to ABSL-3. Following biosafety approval to conduct SARS-CoV-2 experiments under ABSL-2 (IBC approval # P2005190107), subsequent mouse experiments were conducted in ABSL-2 facilities. Mice received *ad libitum* access to standard rodent chow and reverse osmosis H_2_0 in microisolator cages. Mice were acclimated to ABSL-2 or ABSL-3 facilities prior to infections and housing conditions in facilities were maintained under ∼14:10 light dark cycles (14.5 h lights on 06:30 h EST; 20:00 h EST lights off 9.5 h) (**Supplemental Table 3**). All animal procedures were approved by the Johns Hopkins University Animal Care and Use Committee (MO22H358; MO25H376).

### SARS-CoV-2 infections and monitoring

Experimental mice were anesthetized using a ketamine/xylazine cocktail (80 mg/kg ketamine, 5 mg/kg xylazine) administered by intraperitoneal (IP) injection prior to intranasal (IN) infection with either 10^6^ TCID_50_ of the mu variant of SARS-CoV-2 (B.1.621), that contains a naturally occurring N501Y mutation in the spike protein conferring natural viral replication in mice^104^ that was resuspended in DMEM media for a total volume of 30 µm. Mock infections were performed using 30 µm of media (DMEM). Mice were monitored daily for 7 days and weekly thereafter for body mass, rectal temperatures, and clinical signs of disease (i.e., hunched posture, piloerection, dyspnea, and/or an absence of escape response and were assigned a score of 1 per sign for a total scale of 0-4). Cumulative clinical disease scores were totaled across 7 dpi for individual experimental mice to reflect disease severity.

### Tissue collection and serum

Subsets of experimental mice were administered with lethal doses of ketamine/xylazine (160 mg/kg ketamine, 10 mg/kg xylazine) and were euthanized at 2, 4, 7, 42, or 84 dpi, followed by intracardiac bleeds for exsanguination. Blood was collected into heparinized tubes for serum separation and was centrifuged at 10.0 × g for 30 minutes at 4°C and storage at -80°C. At each time point, tissues were collected from mock-infected prior to SARS-CoV-2 infected mice, and all infected or mock-infected cages were collected in alternating orders between sexes. Tissue was collected from extrapulmonary sites (e.g., gonads, WAT, spleen, small intestines (∼0.5 cm of duodenum), brain, and olfactory bulb) prior to collecting respiratory tissues, including nasal turbinates, trachea, and lungs that were separated by lobe (cranial, middle, caudal, accessory) and were flash frozen on dry ice for downstream analyses. The left lung lobe was inflated and fixed in 10% zinc (Zn)-buffered formalin, and in a subset of experimental mice, whole head tissue with intact olfactory epithelium were fixed for at least 72 h for histology. Following administration of lethal ketamine xylazine and intracardiac bleeds, a subset of mice were intracardially perfused with 1X PBS to clear blood, followed by ice cold 4% paraformaldehyde to fix brains for staining. After excision, whole brains were stored in 4% paraformaldehyde for 72 hours, then were switched to 15% and 30% sucrose to cryoprotect tissues for at least 24-48 hours until tissues sank and stored for sectioning and downstream staining.

### Infectious virus quantification

Infectious virus titers in tissue homogenates were determined by a tissue culture infectious dose 50 (TCID_50_) assay^105^. Frozen tissues were weighed and homogenized in 500 ul of DMEM with 100U/ml of penicillin and 100 μg/ml (10% wt/vol) in lysing matrix D bead tubes. Infectious virus titers in tissue homogenates were determined by TCID_50_ assay through serial dilutions in 96-well plates confluent with Vero-E6-TMPRSS2 cells incubated at 37°C for 6 days with 6 replicates. Following incubation, cells were fixed with 10% neutral buffered formalin overnight. Plates were then stained with naphthol blue for visualization and infectious virus titers were determined through the Reed and Muench method. Virus titers were normalized by volume and weight.

### RNA extractions and PCR (qPCR and ddPCR)

For RNA processed for digital droplet PCR (ddPCR), frozen tissues were weighed and homogenized in 1 mL of cold TRIzol in lysing matrix tubes to extract RNA according to manufacturer’s instructions. Genomic viral RNA (gRNA) and subgenomic viral RNA (sgRNA) used a common upstream leader sequence in the 5′ untranslated region (UTR) with separate reverse primers to amplify gRNA or N-sgRNA. 5’ to 3’ gRNA probe (TGTCACTCGGCTGCATGCTTAGT) and sgRNA probe (ACGTTTGGTGGACCCTCAGATTCA) were used with reverse gRNA primers (GCAGCCTGCAGAAGATAGA) and sgRNA primers (CCACTGCGTTCTCCATTC) followed by droplet generation and reading according to the manufacturer’s protocols using the QX200 Droplet Digital PCR System. GADPH was used as a housekeeping gene (5’-GTGGAGTCATACTGGAACATGTAG-3’, 5’-AATGGTGAAGGTCGGTGTG-3’) and copies per well were log10 transformed and normalized to tissue weight. Reverse primers were capable of binding within the N regions within gRNA, however, longer amplicons (>2000 nucleotides) did not have time to amplify due to their large size^106^.

For quantitative PCR (qPCR), up to 1 x 10^6^ cells were isolated from lungs using the EasySep Mouse F4/80 positive selection kit. Single cell suspensions were generated from lung tissue after enzymatic digestion in collagenase and DNAse, followed by mechanically dissociating any remaining tissue and filtering cell suspensions using a 70-μm filter. Red blood cell (RBC) lysis was performed in cell suspensions using ACK lysis buffer then were washed and counted using an automated cell counter (Nexcelom) and viability was determined with AO/PI. RNA was isolated from cells and tissue stored at -80°C using an RNeasy Plus universal mini kit. Frozen tissues processed for qPCR were weighed and homogenized in 1 mL of Qiazol in lysing matrix tubes and RNA was extracted according to the manufacturer’s protocol using an RNeasy kit (Qiagen). RNA was quantified using a NanoDrop, concentrations were normalized between samples, and 250 μg was used to synthesize cDNA using High-Capacity cDNA Reverse Transcription Kit. 1 μl of undiluted cDNA was directly used in a Taqman Real-Time PCR assay in a 96-well plate, each sample was plated with two technical replicates per target. Taqman probes (*Xist*: Mm01232884_m1, *Kdm6a*: Mm00801998_m1, *Tlr7*: Mm00446590_m1, *Irf7*: Mm00516793_g1) were received from ThermoFisher from the Taqman probe catalog (4331182). β-actin was used as a housekeeping gene and endogenous control (Applied Biosystems 4352933E) to normalized expression between samples for each probe, and samples were normalized to wildtype XX females for each gene.

### Multiplex Luminex cytokine and chemokine assay

Cytokine and chemokine concentrations in lung and brain homogenates (inactivated with 0.5% Triton X-100) were measured in duplicate using the ProcartaPlex Mouse Immune Monitoring Panel, 48plex (Thermofisher EPX480 20834-901) following the manufacturer’s instructions and samples were acquired on the xMAP Intelliflex system with a 30 µl acquisition volume, DD gate ranging from 7000 to 17000, and standard reporter gain setting and a minimum bead count of 50. The data was exported as a .csv format and data was analyzed using the ProcartaPlex Analysis App on ThermoFisher Connect. Concentrations of samples were calculated by plotting the expected concentration of the standards against net MFI using a 4 PL curve fit. Undetectable values were set to 50% of the lower limit of detection according to the manufacturer.

### Flow cytometry

Single cell suspensions were generated from spleens or from dissected hippocampi tissues kept on ice in HBSS after collection, by gently dissociating tissue with a plunger in ice cold FACS buffer and transferring the solution to a 70-μm filter to mechanically dissociating any remaining tissue and filtering cell suspensions. For brains, immune cells were separated by slowly and gently layering cell homogenates suspended in 30% percoll on top of a 70% percoll layer to create a percoll density gradient (30/70%) followed by centrifugation for 30 min (800 g, acceleration 2, no brake) at room temperature to separate cells from the myelin and debris. Cells were collected at the interface of the 30 and 70% percoll^107,108^. RBC lysis was performed in cell suspensions using ACK lysis buffer then were washed and resuspended in 1 mL of 1X PBS (protein-free media) for viability staining and counting. Cells were counted using an automated cell counter (Nexcelom) and live cell numbers and viability was determined with AO/PI. 1 x 10^6^ live cells were used for each sample and cells were stained for viability in 1X PBS, followed by Fc receptor block (anti-CD16/32) for non-specific antibody staining. Cells were stained in antibody cocktails for 30 min in the dark, washed, and cells were fixed in 4% paraformaldehyde. For panels using intracellular staining, cells were fixed and permeabilized using the Foxp3/transcription factor staining buffer for intranuclear proteins or Cytofix/Cytoperm for cytokines. Cells were stained with the intracellular antibody cocktail in FACs buffer, then fixed in Foxp3/transcription factor staining buffer or Cytofix/Cytoperm. Compensation was performed with cells and OneComp eBead compensation beads, and cells were acquired using the spectral flow cytometer (Cytek Northern Lights). Cell populations were unmixed with autofluorescence extraction and were gated based on unstained and FMO controls and all data was analyzed using FlowJo v.10 software. Additional details for antibodies used in flow cytometry panels and cell markers used to identify populations are listed in **Supplemental Table 4 and 5**.

Gating strategies for the Myeloid flow cytometry panel are in **Extended Data Fig 5A**. Events were first gated for single cells and red blood cell (RBC) exclusion. Viable CD45+ cells were gated on CD11b expression followed by Ly6C and Ly6G to identify myeloid immune cell subsets, including monocytes (Ly6C high+Ly6G-), neutrophils (Ly6C+Ly6G+), and macrophages (Ly6C low-med expression, Ly6G- followed by F4/80) followed by assessing subpopulations based on activation and cell surface expression. CD11b negative populations were assessed for MHC II and CD11c expression, followed by CD8 and CD11c to identify conventional dendritic cells (cDC1s) and negative cells were gated on Ly6C and B220 to identify plasmacytoid DCs (pDCs). Antibodies used for spectral flow cytometry include a pre-titrated base panel from the Cytek Tonobo Mouse TBNK/Myeloid/Treg kit including: violetFluor 450 Ly6G, violFluor500 CD45, Ghost Dye Violet 540 – viability, FITC CD8, PE F4/80, PerCP-Cy5.5 Ly6C, PE-Cy5 CD3, Pe-Cy7 CD11c, APC CD11b, redFluor 710 CD45R (B220), APC-Cy7 IA/IE (MHC II), with additional markers including BV605 CD86, BV785, CD192 (CCR2).

Gating strategies for Lymphoid panel 1 are in **Extended Data Fig. 6A**. Events were gated for single cells and RBC exclusion. Viable CD45+ cells were gated on CD3 followed by CD4 to identify CD4+ T cells (CD4+CD8-) and CD8 expression to identify CD8+ T cells (CD8+CD4-). CD4+ cells were further assessed for co-expression of CD25 and intracellular staining of Foxp3 to identify T regulatory cells (Tregs) (CD25+Foxp3+) subsets and T helper (Th) cells (Foxp3-). Th and CD9+ populations were further categorized by immune checkpoint expression through CD223 (Lag-3), CD279 (PD-1), cell surface markers, or intracellular expression of CD152 (CTLA-4), with cell activation quantified by CD69. Antibodies used for spectral flow cytometry include a pre-titrated base panel from the Cytek Tonobo Mouse TBNK/Myeloid/Treg kit including: violetFluor 450 CD69, violFluor500 CD45, Ghost Dye Violet 540 – viability, cFluor V610 CD4, FITC CD8, PE Foxp3, PerCP-Cy5 CD152, PE-Cy7 CD279, APC CD25, CD223 redFluor 710, APC-Cy7 CD3.

Gating strategies for Lymphoid panel 2 are in **Extended Data Fig. 6B**. Events were gated for single cells and RBC exclusion. Viable CD45+ cells were gated on CD49b and NK1.1 expression followed by CD3 to identify lymphoid immune cell subsets. Cells expressing NK1.1 and CD49b were assessed for CD45R (B220) or CD3 to identify natural killer (NK) cells (CD3-) and NKT cells (CD3+B220-). Cells expressing CD3 were further assessed for TCRβ expression for total T cells (CD49b-NK1.1-CD3+ TCRβ+), followed by CD4 or CD8. CD4+ (CD4+CD8-) and CD8+ T cells (CD8+CD4-) were then categorized by CD62L and CD44 expression to assess naïve (CD62L+CD44-), central memory (CD62L+CD44+), and effector memory (CD62L-CD44+) T cell populations. CD3 negative populations were assessed for B220 and CD19 expression to identify B cells. Antibodies used for spectral flow cytometry include a pre-titrated base panel from the Cytek Tonobo Mouse TBNK/Myeloid/Treg kit including: violetFluor 450 NK1.1, violFluor500 CD45, Ghost Dye Violet 540 – viability, cFluor V610 CD4, FITC CD8, PE CD62L, PerCP-Cy5.5 TCRb, PE-Cy5 CD19, Pe-Cy7 CD44, APC CD49b, redFluor 710 CD45R (B220), APC-Cy7 CD3.

### Pulmonary histopathology

Fixed left lung lobes were embedded in paraffin and stained with hematoxylin and eosin (H&E) solution. Slides were digitally scanned using an Olympus V200 research slide scanner and microscopically examined with computer-assist Olyvia software. Pathological lung lesions were evaluated for apoptosis/necrosis, alveolar fibrin, acute inflammation, congestion, lymphoid aggregates, AT2 hyperplasia, and mucous cell metaplasia and were assigned a semi-quantitative histopathology severity score by percent of total tissue affected on each sample section on a 0-5 scale (0 = no lung lesions, 0% affected, 1 = minimal, 1% to less than 10%, 2 = 10% to less than 25%, 3 = moderate, 25% to less than 50%, 4 = marked, 50% to less than 75%, 5 = severe, 75% to 100% affected). Histopathological scoring was performed by a board-certified veterinary pathologist blinded to treatment groups, sex, and outcomes, to measure both severity of inflammation and the extent of inflammation across two separate sections of lung per mouse.

### Immunofluorescence staining

Thirty µm coronal sections were prepared from fixed brain tissue using the Microm Cryostat into 4 serial sections in 1X PBS and was stored in cryoprotectant at -80°C until use. Ionized calcium-binding adaptor molecule 1 (Iba1) protein was selected for staining due to its specificity to macrophages (including microglia) in brain parenchyma and is constitutively active that will label the entire cell body including processes to allow for detailed morphological assessment^109^ and was double-labeled with TMEM119. Free-floating brain sections were washed and antigen epitope retrieval was performed using 1X citrate buffer. Sections were blocked for 1 h (2.5% normal goat serum (NGS), 0.2% Triton X-100, in 1X PBS) at room temperature, followed by an overnight incubation with gentle agitation (∼185 rpm) in primary antibody solution with (1:1000 anti-Iba1 rabbit monoclonal), (1:1000 anti-Iba1 chicken polyclonal; 1:500 anti-TMEM119 rabbit polyclonal), or (1:500 anti-CD3 rat monoclonal; 1:500 anti-Ly6G rabbit monoclonal). The next day, sections were washed then incubated with respective secondary antibodies (1:1000 goat anti-rabbit Alexa Fluor 488, 1:1000 goat anti-rabbit Alexa Fluor 555, 1:1000 goat anti-chicken Alexa Fluor 488, or 1:1000 goat anti-rat Alexa Fluor 488; in 2% NGS; 0.2% Triton X-100, in 1X PBS) for 2 h at room temperature. Sections were washed, mounted on slides, dried, and cover slipped with hard set Vectashield antifade mounting medium with DAPI.

Additional free-floating brain sections were stained with either NeuN and DCX, or ZO-1 primary antibodies. To quench autofluorescence, free-floating sections were initially incubated for 30 minutes in 0.01% Sudan Black B in 70% EtOH solution and then rinsed in 70% EtOH solution for 5 minutes, followed by a final rinse in PBS for 5 minutes. Sections were then blocked for 1 h (5% NGS in 1X PBS) at room temperature, then incubated overnight with gentle agitation in primary antibody solution (1:200 anti- NeuN mouse polyclonal and 1:250 anti-DCX rabbit polyclonal co-stain; or 1:200 anti- ZO-1 rabbit monoclonal) at 4°C. The next day, sections were washed then incubated with secondary antibody (1:2000 AF568 goat anti-mouse IgG; 1:2000 AF488 goat anti-rabbit IgG; in 1X PBS) for 2 h at room temperature. Sections were washed, mounted on slides, dried, and cover slipped with Vectashield antifade mounting medium with DAPI and sealed with clear nail polish.

### Imaging and morphological characteristic analyses

Immunofluorescence images were acquired using the AxioObserver Z1 on the LEICA Thunder widefield microscope, using the 40 X objective with the DAPI, FITC (488 nm laser), (594 nm laser) and BF filter. For analysis, tile scanned 40X images of the hippocampus were obtained, and maximal projection images from 30 μm Z-stacks with a 1μm interval were used to quantify Iba1+ cell numbers and for morphological classification scoring. Cells were exhaustively counted throughout each section throughout the hippocampus [dentate gyrus (DG), CA3, CA2, and CA1]. For each section examined, the area was calculated from manual tracings and outlines of the hippocampus in ImageJ. Guidelines for manual segmentations were determined using the Allen Institute mouse brain atlas as a reference guide (https://mouse.brain-map.org/static/atlas). Iba1+ cells were classified into 4 main morphological types based on cell shape, branches, and processes. These cell types consisted of: microglial cells with thin, ramified processes (homeostatic), hyper-ramified with thick long processes, thick and stout-rod like bodies, and rounder/ameboid and reactive microglia (activated); terms defined by^110^.

For quantitative assessment of individual cell morphological characteristics, 40X maximum intensity projections of 30μm Z-stack with 1μm step intervals were used to visualize all ramifications and cell processes^111^ and were conducted using FIJI ImageJ software (https://imagej.net/software/fiji/). Individual microglia were selected for individual morphological segment characteristics based on an unbiased approach following inclusion criteria to identify cells (entire soma in view, cell contained within Z-stack and field of view, none or minimal overlap with neighboring microglia for accurate segment tracking, and sufficient signal fluorescent quality to identify ramifications from background). Cells that fit within the inclusion criteria within a full field of view using the Grid Plugin in ImageJ/Fiji by assigning numbers to grid squares containing at least 1 positive-inclusion microglia and were randomly selected with a number generator. In FIJI, image adjustments for cell structure binarization was applied using the same settings across all images. Brightness/contrast was adjusted to a constant value, an unsharp mask filter was performed (pixel radius of 3 and mask weight of 0.6), noise de-specking was performed to eliminate single-pixel background fluorescence, then, the minimum threshold (0-255) was adjusted to a constant value. Images were de-speckled, close binary was applied (up to 2 pixels), and noise outliers were removed (pixel radius of 2 threshold 50). Images were converted to binary files and were adjusted using the paintbrush to isolate the cell of interest. The original unedited image was used as a frame of reference to connect fragment processes ^111^ and isolated cells were saved as separate modified isolated cell files. Binarized isolated cells were then skeletonized using the AnalyzeSkeleton (2D/3D) plugin and sholl analysis was performed using SNT Sholl Analysis plugin for FIJI by generating concentric circles were added to the center of the soma increasing at a 5 μm radius. Fractal analysis was performed by using the binarized isolated cell image, converting to an outline, and using the FracLac plugin for FIJI (Grid design settings Num G to 4, and include convex hull and bounding circle of the cell)^111^.

Immunofluorescence images for NeuN, DCX, ZO-1, and FJC were acquired on the Zeiss AxioScan Z.1 using the 20 X objective with the DAPI, FITC (488nm), and 568nm lasers. For each animal, at least one brain slice and up to 4 slices were stained and analyzed. The region of interest (ROI), the hippocampus, was isolated in each section using QuPath and exported into ImageJ. In ImageJ, a boundary around the outermost edge of the hippocampus was manually drawn using the Allen Institute mouse brain atlas as a reference. The area of the entire hippocampus was calculated from this boundary. Immunofluorescence staining for NeuN and ZO-1 were quantified using positive pixel analysis in ImageJ with the Triangle threshold function applied to isolate percent positive area within the hippocampus. DCX+ cells were manually counted in ImageJ using the cell counter tool and normalized to the area of the ROI.

The DCX+:NeuN ratio was calculated by normalizing the number of DCX+ cells counted to the percent positive area of NeuN stain in the ROI.

### Steroid measurements

Concentrations of sex steroids (e.g., testosterone and estradiol) were measured in hormone-extracted serum samples using diethyl ether at a 1:5 sample-to-ether ratio. Testosterone concentrations were measured using mouse IBL Testosterone Kit and Estradiol was measured using the Milliplex multi-species hormone magnetic bead kit (MSHMAG-21K), samples below LOD were listed as not detectable (n.d.) in graphs.

### Cognitive behavioral assessments

Mice were evaluated for behavior at set timepoints following infection in ABSL-3 facilities, with subsequent experiments conducted in ABSL-2 following biosafety approval. Cognitive behavioral assessments, Full behavioral phenotyping assessments were conducted and are located in **Supplemental Materials** (**Fig. 1**).

### Spontaneous Alternations in a T-maze

Spatial working memory was conducted in a T-maze apparatus constructed to fit and accommodate behavioral testing within the biosafety cabinet. Mice were habituated to the testing room for a minimum of 30 min prior to behavioral testing, and an additional 30 min in a separate biosafety cabinet under nearly identical sound, vibration, and light exposure prior to testing. Following habituation, mice were placed into one arm facing the same direction and were left undisturbed and allowed to freely move through the maze for 5 min. Locomotor activity was recorded and tracked during the testing interval using ANYmaze software, and arm entry patterns were assessed. After completion of the T-maze task, mice were removed from the maze and returned to their home cage. The T-maze apparatus was thoroughly cleaned with Vimoba then 70% EtOH and wiped dry between trials to eliminate residual odors. Arm entries were defined as all four limbs crossing into the arm and percent alternation was calculated using: [(alternation/maximum possible number of alternations) x 100]. The maximum possible alternations were the total arm entries -2.

### Novel Object Recognition (NOR) Task

To assess visual recognition memory, the NOR test was performed to quantify the amount of time spent investigating familiar versus novel objects. The NOR procedure consists of three testing phases, including habituation, familiarization, and the testing trial. During the habituation phase, mice were habituated to the testing environment and freely explored the OF apparatus for 10 min (served as the OFT; methods described in Open Field) and was performed 24 h prior to familiarization and testing session to reduce stress and the potential for a neophobic response. The next day, following habituation to the testing room and biosafety cabinet, two identical objects (A/A) or (B/B) were placed equal distance from the wall of the open field box for the familiarization session. Experimental mice were placed in the OF maze and were given a 10 min-period to investigate and explore their environment. Following the familiarization session, mice were returned to their home cages and left undisturbed for an 4 h inter-session-interval (ISI). During the testing trial, one of the two identical objects were replaced with a novel object (A/B) or (B/A) in the identical location as the familiar object, then were allowed to freely explore their environment for 10 min.

Exploration and locomotor activity and track patterns were recorded and monitored using ANYmaze software across all trials. Object investigation scoring, including the total number of investigations, investigation time (seconds), and distance travelled was analyzed during the familiarization session (10 min session interval) using automated scoring software from recorded video footage (ANYmaze) by tracking mouse head orientation and interaction within a 3 cm investigation zone from objects, and excluding time the animal spends in the center of the object (climbing without active interaction). Automated scoring was conducted to determine baseline object exploration between either object A or B (familiar object), and demonstrate no inherent object preference. Investigation time was manually scored from recorded video footage during the novel object testing trial; investigation was scored until a total of ∼20 sec of exploration time was reached between object A and B^112^. Investigation time was defined as direct interaction and exploration of the object by directing the nose towards the object including actively engaging in sniffing behaviors or interaction with their hands, but excluded behaviors such as standing on the object without direct active interaction. Object preference was reported as a percentage of total investigation time between the familiar and novel object per mouse. For this NOR task, the objects used included a T25 flask or a 50 mL conical tube, and a 1L glass orange bottle lid that contained different textures, colors, and were similar sized objects. Object order and the location of the novel object was randomized across treatment groups and cohorts, and between all testing intervals, the apparatus and testing objects were thoroughly cleaned with Vimoba then 70% EtOH and wiped dry between trials to eliminate residual odors.

### TLR7 agonist and inhibitor treatment

Into an open standard rodent diet with 15% kcal fat (D23100303B), 40 mg/kg of selective TLR7/8 inhibitor Enpatoran (M5049) was incorporated by Research Diets Inc, with batch matched diets used as controls (D11112201), following previous *in vivo* experiments that confirmed brain permeability^67^ and prior testing in experimental mice with autoimmune conditions^97^. Experimental mice were placed on batch-matched control diets for 7 days prior to infection to acclimate to the change in rodent chow. Animals were either maintained on the control diet for the duration of the study or were switched to the TLR7 inhibitor (40 mg/kg) diet after recovery from acute disease at 7 dpi. Research diets were weighed and replaced every 3-4 days; diets were stored at 4°C and were used within <6 months of development.

A TLR7 agonist, imiquimod, was used to stimulate TLR7 by epicutaneously administering 50-51 mg of a 5% topical formulation cream aliquot (NDC 45802-368-62) to the ventral inner surface of the right ear 3 times weekly, or were applied with cream lotion was used as a vehicle control [administered to (Mock - Vehicle; SARS-CoV-2 – Vehicle; SARS-CoV-2 -TLR7 inhibitor)]. Topical administration of ∼50 mg of 5% imiquimod cream (containing ∼2.5 mg of active ingredient) is lower than the established dosage used for autoimmune disease or psoriasis, and utilized a modified dosing frequency to be better tolerated *in vivo* (PMID: 42052298). Experimental mice were treated with either a TLR7 agonist or vehicle control starting after recovery from acute SARS-CoV-2 infection at 7 dpi.

## Statistical Analyses

All statistical analyses were performed in GraphPad prism with statistical tests indicated in figure legends. Statistical significance was determined by an ordinary two-way ANOVA for body mass (AUC) analyses, semi-quantitative histopathological assessments, infectious virus titers, flow cytometry, immunofluorescence quantifications, behavior, steroids, and qt-PCR, repeated measures two-way ANOVAs were used for body mass analyses across time (0-82 dpi), followed by Tukey’s multiple-comparison post-hoc tests. The Kruskal Wallis test was used for cumulative clinical disease scores and viral RNA assessments, followed by the Mann-Whitney test for multiple comparisons. Unpaired individual two-tailed parametric t-tests were used to compare individual cytokine/chemokines between infected male and female lung homogenates at 2 dpi. Linear mixed effects models were used for datasets that sampled multiple cells from the same individual to account for within-subject clustering. Data were considered statistically significant if p<0.05, and are represented as mean ± standard error of the mean, or as the median (cumulative clinical score). For all data, individual data points are shown for individual experimental mice with the exception of microglial segment characteristic analyses which is specified in the figure captions. Experimental conditions, replicates, and exact sample size (n) for each experimental group/conditions is listed in the corresponding source data excel sheet. Statistically significant differences and comparisons between groups, across time, or within a sex were all denoted with different colored connecting solid or stippled lines and were listed in figure captions. Mock-infected male and female mice were always collected at the same timepoints as all SARS-CoV-2 infected mice for all experiments and showed no within-group difference between collections across timepoints, therefore were collapsed for comparisons. Sample size was determined based on previous historical mouse data with other pathogens, that includes similar infected and mock-infected sample sizes and both sexes within respective groups. Sample size was also determined using power calculations based on preliminary data to detect sex differences at 42 dpi within the SARS-CoV-2-infected group for measures including behavior. Sample sizes of 10-15 mice/sex or genotype at an alpha of 0.05 and power of 0.8 were calculated. GraphPad Prism was used to evaluate the contributions of circulating sex steroid concentrations and ChrX dosage on cognitive behaviors in infected XY* mice at 42 dpi. Separate multiple linear regression (MLR) models were fit in gonadal males (XYM, XXYM; testosterone n=19, 7 or 12 per group) and gonadal females (XXF, X0F; estradiol model n=14, 7 per group) so that circulating sex steroids were modeled in gonadal types. Spatial memory assessment scores (correct % of spontaneous alternations) were the dependent variable (lower scores indicate greater impairment) in both models. Predictors were steroid concentrations (continuous), ChrX dosage (2 vs 1 ChrX; 1X as the reference), and “hormone x ChrX dosage” interaction. Steroid concentrations were mean-centered within each model by subtracting the mean steroid level in that model) so ChrX dosage coefficients were the difference between 2 vs 1 ChrX at average steroid concentrations. Steroid coefficient is the slope in 1 ChrX mice. Coefficients are reported as unstandardized β, with 95% confidence intervals (profile-likelihood), and tests used type III sums of squares with two-sided α = 0.05. Multicollinearity was assessed for model stability (testosterone VIF 1.0 - 1.39, estradiol VIF 1.05 - 1.84) and residuals were inspected for normality and homoscedasticity. Overall model fit is reported as F statistic with degrees of freedom and R^2^. For data visualization, individual values were mapped on scatterplots, with unadjusted trendlines for each ChrX dosage group; statistical inference is from the regression models. Model formula for testosterone [(Spontaneous alternation ∼ Intercept + Testosterone (pg/ml + # ChrX + Testosterone (pg/ml : # X Chromosomes)] and estradiol [(Correct % spontaneous alternation ∼ Intercept + Estradiol pg/mL + # X Chromosomes + Estradiol pg/mL : # ChrX)]. Per mouse data and model outputs are available in the source data file.

## Extended Data Figures

**Extended Data Figure 1. Development of mouse model to explore sex differences in acute and post-acute SARS-CoV-2 infection.** To develop a mouse model of sex differences in SARS-CoV-2 pathogenesis, male and female mice were intranasally (IN) infected with of SARS-CoV-2 (B.1.621) at 10, 30, or 80 weeks of age, and monitored mice daily for body mass changes (**A-B**) from baseline (stippled line) with the area under the curve (AUC) calculated for comparisons (**C**) through 7 days post-infection (dpi) in inbred C57BL6/J mice. To confirm sex and age differences in acute SARS-CoV-2 pathogenesis, outbred CD-1 mice were infected at 10, 30, or 80 weeks of age, and monitored mice daily for body mass changes (**D-E**) from baseline (stippled line) with the area under the curve (AUC) calculated for comparisons (**F**) through 7 dpi. Data are represented as mean ± standard error of the mean (SEM) from two to three independent replications (n =4-10/group). Statistical significance was determined by a two-way ANOVA (repeated measures and ordinary ANOVA) followed by Tukey’s multiple-comparison test, Kruskal-Wallis test, or the Mantel-Cox test. Asterisks (*) represent significant differences (p<0.05) between groups; black solid lines reflect sex differences in multiple comparisons tests between males and females following a significant interaction; colored lines represent differences within a sex based on ages of infected mice **(C, F)** (green= males; purple= females). Open circle symbols in the figure represent individual mock-infected mice, closed symbols represent B.1.621 infected mice. Closed circles represent C57BL/6 mice, closed squares represent CD-1 mice, with colors reflecting sex (green = males, purple = females) (**A-F**).

**Extended Data Figure 2. Limited detection of SARS-CoV-2 subgenomic (sg) and genomic (g) RNA in extrapulmonary tissues.** Male and female mice were infected with SARS-CoV-2 (B.1.621) or mock-infected, and euthanized at 7, 42, or 84 days post infection (dpi) for tissue collection. SARS-CoV-2 sg and g RNA was quantified in extrapulmonary tissues, including brain (**A,D,H**), olfactory bulb (**B,E,I**), white adipose (**C,F,J**), and small intestines (**G,K**). Interleaved whisker plots for infected male and female mice are represented by closed bars, with individual points representing individual mice. Data are represented as mean ± SEM from one to two independent replications (digital droplet PCR n=4-9/group). Statistical significance was determined by a two-way ANOVA followed by Tukey’s multiple-comparison test or Kruskal Wallis test followed by Mann Whitney multiple-comparisons test. Open circles/bars represent mock-infected mice and closed circles/bars represent B.1.621 infected mice, with colors reflecting sex (green=males, purple=females).

**Extended Data Fig. 3. Extended cognitive behavioral tests, microglial segment characteristics in the hippocampus, and microglia flow cytometry gating strategy.** Measures of spatial working memory were conducted in mice at 7 dpi by measuring the correct percentage of spontaneous alternations (**A**) and number of arm entries (**B**) in the T maze. The total distance travelled during the T-maze task was measured (**C**). Novel object preference (NOR) was quantified at 84 dpi (**D**). Mock-infected control male and female mice were assessed at each timepoint and during the same time interval as SARS-CoV-2 infected mice. Because there was no significant differences among mock-infected mice based on dpi, mock-infected males and females were collapsed across time for statistical comparisons. SARS-CoV-2 infected and mock-infected male and female mice were euthanized at 42 or 84 dpi, for collection of either fresh brain tissue for flow cytometry or perfused brain tissue for immunofluorescence staining with Iba1 co-stained with TMEM119 (**E**), or were stained with Ly6G and CD3 **(F)**. Microglial morphological states (hyper-ramified and rod-like) were reported as percentages of total microglia, and were classified from maximal projected 30 μm z-stack images of microglia in the hippocampus with intermediary classification phenotypes (hyper-ramified, rod-like) (**G,H**). Morphological parameters and segment characteristics of individual cells were assessed from immunofluorescence images for Iba1 using 40X maximum intensity projection 30 μm z-stack images (1 μm stack interval) to visualize processes in hippocampal microglia. Individual cells from immunofluorescence images were isolated, underwent threshold analysis, were skeletonized for branch information, and outlined for fractional dimension and scholl analysis to quantitatively measure morphological changes. An illustrative schematic of Z-stack projection and morphological analysis pipeline (**I**). Span ratio (**J**), fractional dimension (**K**), circularity (**L**), and lacunarity (**M**) were quantified as additional measures of microglial activation. Individual microglial segment characteristic analysis were randomly isolated with an n=10-20 microglia/mouse performed across 4-5 40X maximal projected Z-stack images in the dentate gyrus, with individual points representing individual microglia. Data are represented as mean ± SEM from two to three independent replications (n =7-8/group). Statistical significance was determined by a two-way ANOVA followed by Tukey’s multiple-comparison test. Asterisks (*) represent significant differences (p<0.05) between groups, with black solid lines reflecting sex differences in multiple comparisons tests between males and females following a significant interaction, and colored solid lines representing differences within sex comparing mock and/or infected mice at different dpi (green=males, purple=females). Open circles/bars represent mock-infected mice and closed circles/bars represent B.1.621 infected mice, with colors reflecting sex (green=males, purple=females).

**Extended Data Fig. 4. Myeloid gating strategy in the brain and impact of SARS-CoV-2 infection on neurogenesis, neurodegeneration, and blood brain barrier (BBB) permeability.** From fresh hippocampal tissue, single cell suspensions were generated by mechanical dissociation followed by a percoll density gradient (30% and 70%) and centrifugation to isolate immune cells from brain tissues. The gating strategy for the myeloid flow cytometry panel used a modified pre-established base (Tonobo) panel from CytekBio (**A**). Cell populations were gated based on unstained and FMO controls. Data acquisition was conducted using an Cytek Northern Lights spectral flow cytometer, and flow cytometry data were evaluated using FlowJo v10. Mature neurons were labeled with NeuN and immature neurons and progenitor cells were labeled with Doublecortin (DCX) in the hippocampus using immunofluorescence images of brain sections in subsets of SARS-CoV-2 and mock-infected male and female mice at 7, 42, or 84 days post infection (dpi; **B**). The percentage of positive NeuN (**C**) or DCX (**D**) staining (% area) was calculated. NeuN:DCX ratios were quantified to assess the relative proportion of mature to immature neurons/neuroblasts within the hippocampus as a measure of adult neurogenesis (**E**). Zonula Occludens-1 (ZO-1) was used to measure tight junctions as an indicator of BBB integrity. Representative images of ZO-1+ staining from immunofluorescence in the dentate gyrus of the hippocampus (**F**), with the % of positive ZO-1 in the hippocampus quantified (**G**). Data are represented as mean ± SEM from two to three independent replications (n =4-8/group). Statistical significance was determined by a two-way ANOVA followed by Tukey’s multiple-comparison test. Asterisks (*) represent significant differences (p<0.05) between groups, with black solid lines reflecting sex differences in multiple comparisons tests following a significant interaction, and colored solid lines representing differences within sex comparing mock and/or infected mice at different dpi (green=males, purple=females). Open circles/bars represent mock-infected mice and closed circles/bars represent B.1.621 infected mice, with colors reflecting sex (green=males, purple=females).

**Extended Data Fig. 5. Gating strategy for myeloid flow cytometry panels and extended flow cytometry cell populations in lungs and spleens.** The gating strategy for the myeloid flow cytometry panel used to identify single cells from lungs or spleens of male and female mice infected with SARS-CoV-2 or mock infected at 4, 7, or 42 days post infection (dpi; **A**). Macrophages were identified as live CD45+CD3-CD11b+Ly6G-Ly6C (low-med), F4/80+ with subsets of activated macrophages further gated on CD86 expression, Monocytes were identified as live CD45+CD3-CD11b+Ly6G-Ly6C (high), with inflammatory monocytes identified by CCR2 expression. For this panel, myeloid cell populations were quantified including CD86+F4/80+macrophages CD206+F4/80+ M2 macrophages, Ly6G+Ly6C+ neutrophils, and pDCA1+ pDCs (**A-H**). Cell populations were gated based on unstained and FMO controls. Representative contour plots show the gating strategy used to identify myeloid cells from single cell suspensions generated from lungs and spleens. Pulmonary immune cells were quantified, including reporting the frequency of CD86+ cells reported out of total macrophages (**B**), along with monocytes (**C**), and neutrophils (**D**), reported as a percentage of total CD45+ cells when infectious virus is present (4 dpi) and post-acute (42 dpi). Frequencies of splenic immune cell populations from the myeloid panel collected at 7 or 42 dpi include F4/80 macrophages (**E**), MHC II+ macrophages (**F**), CD206+ macrophages (**G**), and neutrophils (**H**), reported as percentage of total CD45+ live cells. Data acquisition was conducted using an Cytek Northern Lights spectral flow cytometer, and flow cytometry data were evaluated using FlowJo v10. Data are represented as mean ± SEM from two to three independent replications (n =7-8/group). Statistical significance was determined by a two-way ANOVA followed by Tukey’s multiple-comparison test. Asterisks (*) represent significant differences (p<0.05) between groups, with black solid lines reflecting sex differences in multiple comparisons tests following a significant interaction, and colored solid lines representing differences within sex comparing mock and/or infected mice at different dpi (green=males, purple=females). Open circle symbols in the figure represent mock-infected mice and closed circle symbols with filled bars represent B.1.621 infected mice, with colors reflecting sex (green=males, purple=females).

**Extended Data Fig 6. Gating strategy for lymphoid flow cytometry panels and extended flow cytometry cell populations.** Representative contour plots show the gating strategy used to identify lymphoid populations for lymphoid panels used for flow cytometry, that was generated from single cell suspensions prepared from lungs and spleens collected from male and female mice infected with SARS-CoV-2 or mock infected at 4, 7 or 42 days post infection (dpi) using a Treg panel (lymphoid panel 1) and TBNK panel (lymphoid panel 2) from CytekBio. T cells were identified as live CD45+CD3+ cells, followed by CD4 to identify CD4+ T cells (CD4+CD8-) and CD8 expression to identify CD8+ T cells (CD8+CD4-). CD4+ cells were further assessed for intracellular staining of Foxp3 to identify T regulatory cells (Tregs) and T helper (Th) cells were identified as Foxp3-. Th and Treg cells were further categorized by immune checkpoint expression through CD223 (Lag-3), CD279 (PD-1), cell surface markers, or intracellular expression of CD152 (CTLA-4), with cell activation quantified by CD69 **(A).** Frequencies of CD4 and CD8 cells were assessed for expression of CD44 and CD62L to quantify central and effector memory. Additional lymphoid subsets were quantified by expression of NK1.1 for NK cells, and B cells were identified as CD3- and by expression of CD19 and B220 (**B**). Data acquisition was conducted using an Cytek Northern Lights spectral flow cytometer, and flow cytometry data were evaluated using FlowJo v10. Data are represented as mean ± SEM from two to three independent replications (n =7-8/group). Statistical significance was determined by a two-way ANOVA followed by Tukey’s multiple-comparison test. Asterisks (*) represent significant differences (p<0.05) between groups, with black solid lines reflecting sex differences in the interaction followed by multiple comparisons tests between males and females, and colored solid lines representing differences within sex comparing mock and/or infected mice at different dpi (green=males, purple=females). Open circles/bars represent mock-infected mice and closed circles/bars represent B.1.621 infected mice, with colors reflecting sex (green=males, purple=females).

**Extended Data Fig 7. Gating strategy for lymphoid flow cytometry panels and extended flow cytometry cell populations.** Lung and splenic immune cell frequencies were quantified based on the gating strategy in Extended Data Fig. 6, with NK cells (**A**), NKT cells (**B**), CD4+ T cells (**C**), and the expression of exhaustion markers with or without activation (CD69) including CD279, CD152, and CD223 in CD4+ Th cells or CD8+ T cells (**A-O),** and B cells (**P**), and reported as a percentage of total CD45+ cells. Lymphoid subsets were further gated based on CD44 and CD62L expression for naïve, effector, and central memory (**Q-T**). Data acquisition was conducted using an Cytek Northern Lights spectral flow cytometer, and flow cytometry data were evaluated using FlowJo v10. Data are represented as mean ± SEM from two to three independent replications (n =7-8/group). Statistical significance was determined by a two-way ANOVA followed by Tukey’s multiple-comparison test. Asterisks (*) represent significant differences (p<0.05) between groups, with black solid lines reflecting sex differences in the interaction followed by multiple comparisons tests between males and females, and colored solid lines representing differences within sex comparing mock and/or infected mice at different dpi (green=males, purple=females). Open circles/bars represent mock-infected mice and closed circles/bars represent B.1.621 infected mice, with colors reflecting sex (green=males, purple=females).

**Extended Data Fig. 8. Immunophenotyping and additional behavioral outcomes following SARS-CoV-2 infection of XY* mice.** Adult XY* mice (C57BL6/J background), including gonadal females (XX or X0 chromosome complement) and gonadal males (XY or XXY chromosome complement), were infected with SARS-CoV-2 (B.1.621) or mock-infected at 30 weeks of age. Illustrative schematic diagrams components of sex chromosomes in the XY* mouse model, and a summary table of XY* genotypes **(A)**. Mock-infected XY* mice were monitored for body mass changes over the course of the experiments (**B**), with area under the curve (AUC) calculations showing no changes in body mass in mock-infected mice compared to baseline (stippled line; **C**). Subsets of XY* mice infected with SARS-CoV-2 (B.1.621) were euthanized and infectious virus titers as measured by TCID_50_ in trachea **(D)** and nasal turbinates **(E)**. The dashed line indicates the limit of detection (LOD). Frequencies of F4/80+ (**F**) and MHCII+ (**G**) macrophage populations reported as a percentage of total CD45+ live cells were generated from single cell suspensions prepared from spleens collected from mock-infected and B.1.621 infected XY* mice at 42 days post infection (dpi). Additional behavioral data from mock-infected XY* mice, including percentages of correct spontaneous alternation (**H**) and total number of arm entries (**I**) in the T maze and object recognition preference (**J**) during the novel object recognition task. Scatterplots of spatial cognitive memory scores (correct % spontaneous alternation) in T maze against circulating sex steroid concentrations for multiple linear regression analyses in infected XY* mice. Circulating testosterone (ng/ml) in gonadal males (**K;** XYM, XXYM n=19, 7 or 12 per group) and estradiol (pg/ml) concentrations in gonadal females (**L;** XXF, X0F n=14, 7 per group) at 42 dpi are separately graphed with unadjusted simple linear fit trendlines for each X chromosome group (2 vs 1 ChrX) shown for visualization. Statistical inference is from multiple linear regression models. Data are represented as mean ± SEM from two to three independent experiments (morbidity n=7-13/group, behavior n=7-10/group). Statistical significance was determined by a two-way ANOVA (ordinary and repeated measures) followed by Tukey’s multiple-comparison test. Asterisks (*) represent significant differences (p<0.05) between groups, with black solid lines reflecting sex differences in the interaction followed by multiple comparisons tests between males and females, colored solid lines representing differences within a gonadal sex (1 vs. 2 chromosomes), and colored stippled lines representing differences within sex comparing mock and infected mice (green= males; purple= females). Individual data points represent separate experimental mice with triangle symbols representing mice with one X chromosome, square symbols representing mice with two X chromosomes, and color reflecting gonadal sex (green = males, purples = females). Open symbols represent mock-infected mice and closed symbols represent B.1.621 infected mice.

**Extended Data Fig. 9. Morbidity, immunophenotyping, and additional behavioral outcomes following manipulation of TLR7 after acute SARS-CoV-2 infection**. Adult male and female C57BL6/J mice were infected with SARS-CoV-2 (B.1.621) or mock-infected at 30 weeks of age. Daily morbidity was recorded and shown by the percent change of body mass from baseline (stippled line) through 7 days post infection (dpi; **A-B**). Body mass curves were converted into area under the curve (AUC) values (arbitrary units) for individual mice, where lower AUC values correspond to greater mass loss post infection (**B**). After 7 dpi, male and female mice received either TLR7 inhibitors (M5049) 40 mg/kg in standard rodent chow, topical TLR7 agonist (SARS-CoV-2 TLR7 inhibitor) administered 3 times weekly, or were maintained on batch-matched control diets for 35 days with behavior and experimental collection at 42 dpi. Weekly measures of body mass were reported as the percentage change of body mass from baseline (**C**). RNA was isolated from lung homogenates at 42 dpi for qR-PCR analyses of *Irf7* (**D**), with relative expression normalized to expression in uninfected XX females, with β-actin as a housekeeping control. Single cell suspensions were prepared from the lungs and analyzed by spectral flow cytometry at 42 dpi. Frequencies of F4/80 macrophages in the lungs as a percentage of live CD45+ cells (E), CD86+ macrophages reported as a percentage of total macrophages **(F)**, and neutrophils **(G)** reported as the percentage of total live CD45+ cells were measured. Cognitive function and spatial working memory were assessed by measuring spontaneous alternations in a T maze, with the total number of arm entries (**H**) and novel object preference indexes (**J**). Statistical significance was determined by a two-way ANOVA (ordinary and repeated measures) followed by Tukey’s multiple-comparison test. Asterisks (*) represent significant differences (p<0.05) between groups, with black solid lines reflecting sex differences in the interaction followed by multiple comparisons tests between males and females, and colored solid lines representing differences within sex comparing mock and/or infected mice in different drug treatment groups (green=males, purple=females).

## Source Data

Statistics source data is provided in Excel format, one file for each relevant figure, with the linked figure noted directly in the file.

## Supplementary Information

**Table S1.** Concentrations (pg/ml) of cytokines and chemokines in lungs collected from mock-infected or SARS-CoV-2 infected males and females.

**Table S2.** Concentrations (pg/ml) of cytokines and chemokines in brains collected from mock-infected or SARS-CoV-2 infected males and females.

**Supplemental Fig. 1 and Supplemental Methods.** Extended behavioral tests in SARS-CoV-2 (B.1.621) infected mice.

**Supplemental Table 3.** Reporting animal research. list of experimental animals, strains, and viruses used in this study.

**Supplemental Table 4.** List of antibodies used for flow cytometry and immunofluorescence

**Supplemental Table 5.** Cell markers to identify immune cell populations.

