## Supplementary figures and images for "Mechanisms of sex differences in acute and long COVID sequelae in mice"

### Supplemental Fig 1-9

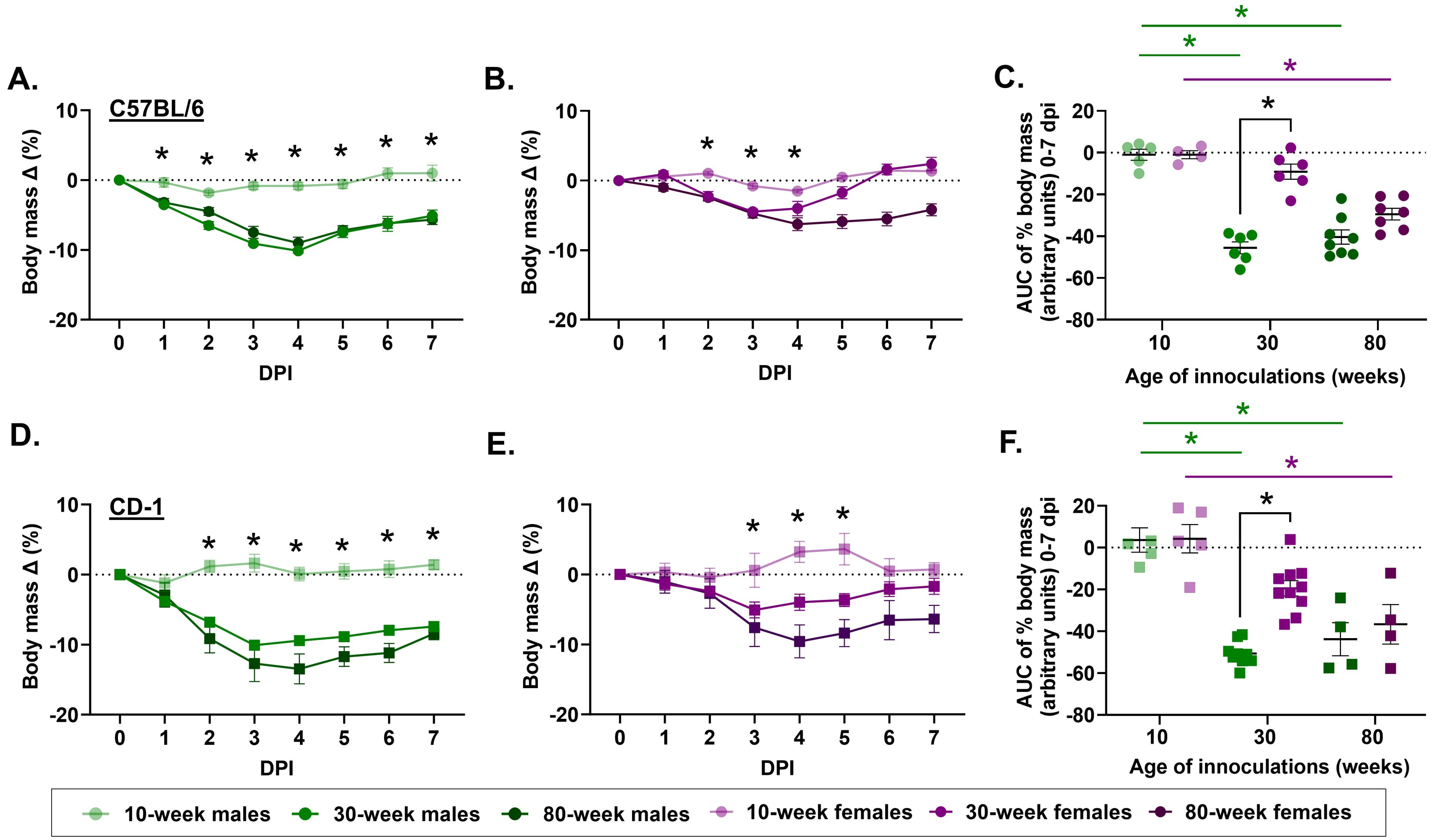

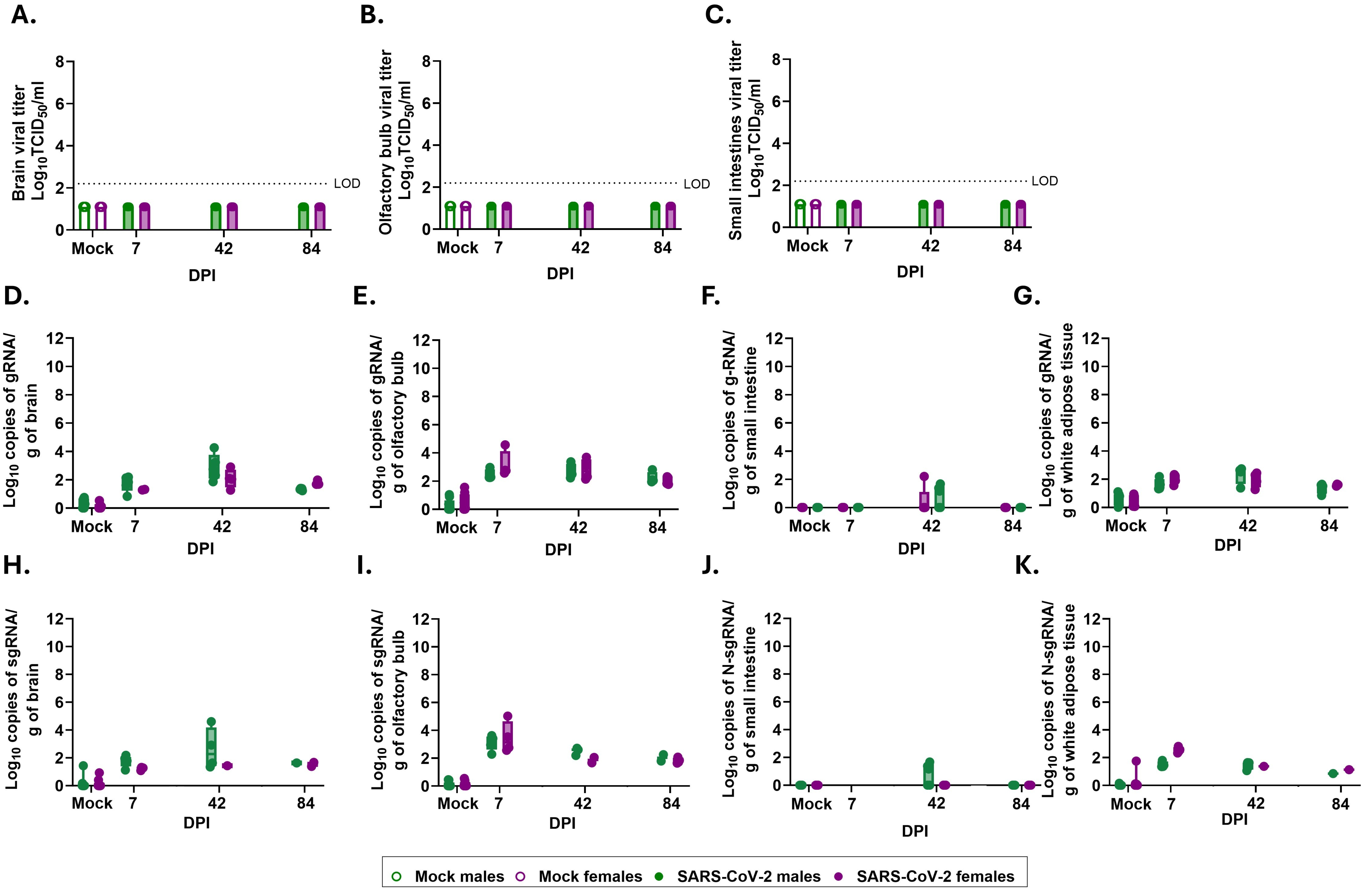

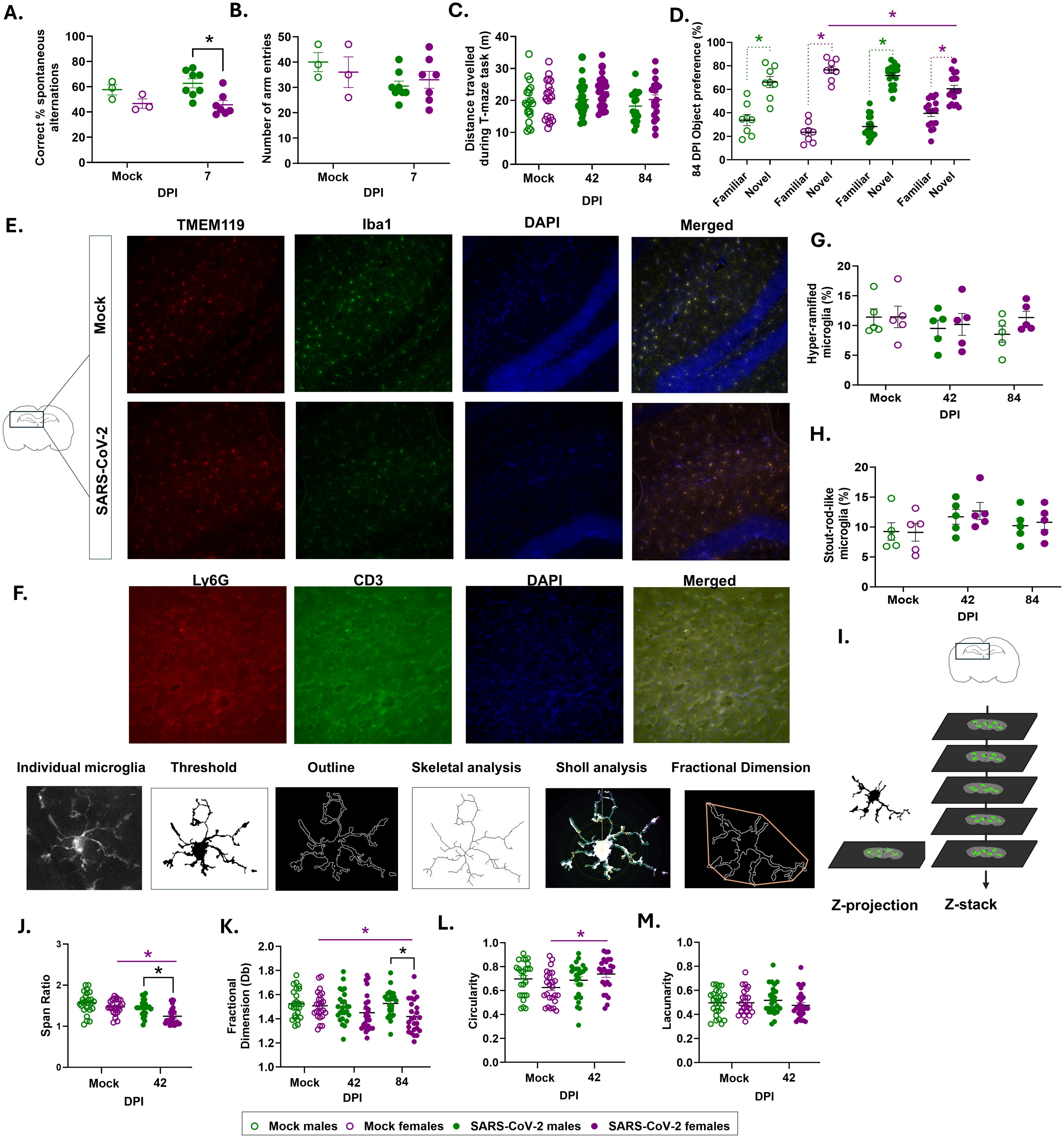

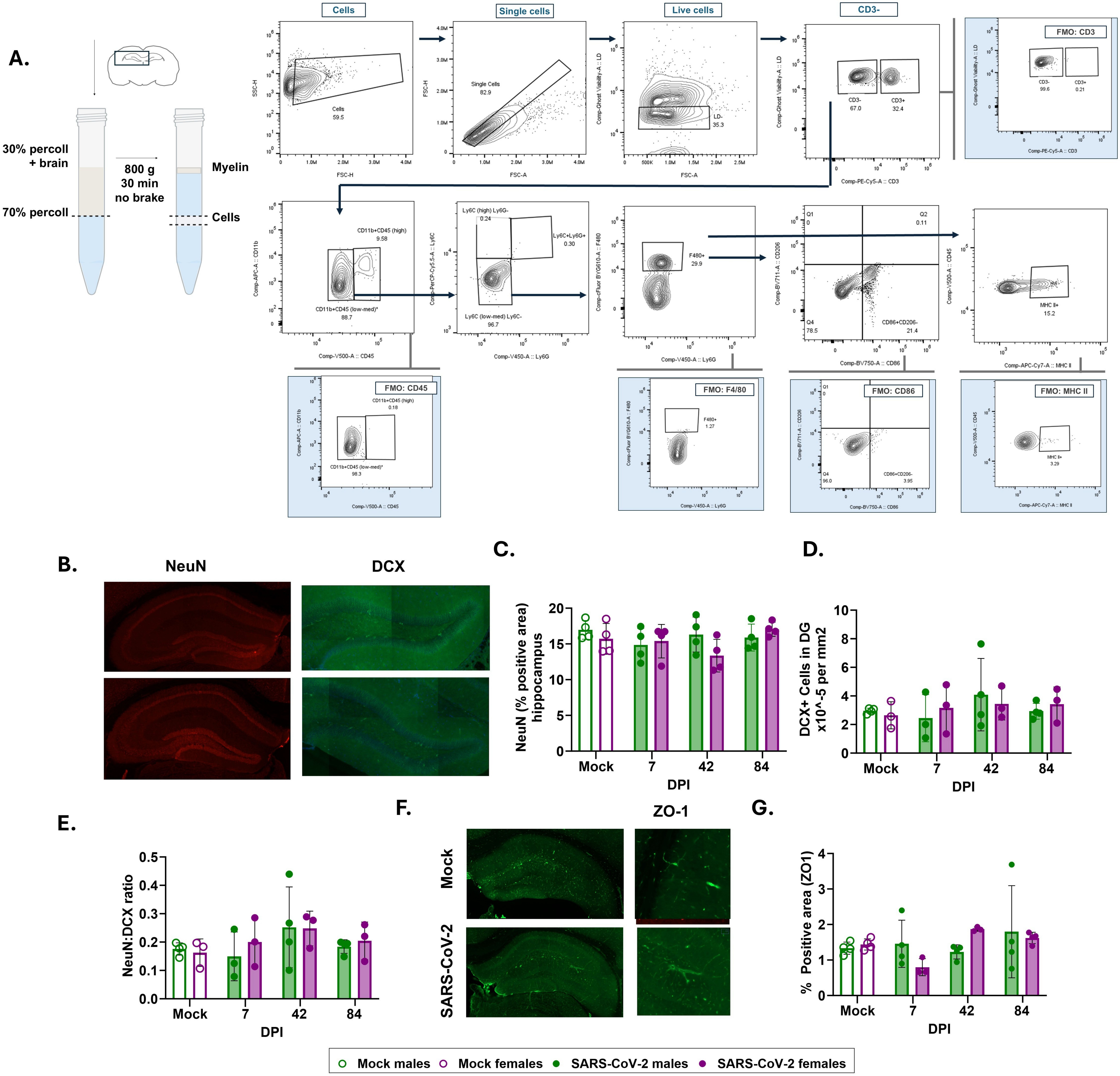

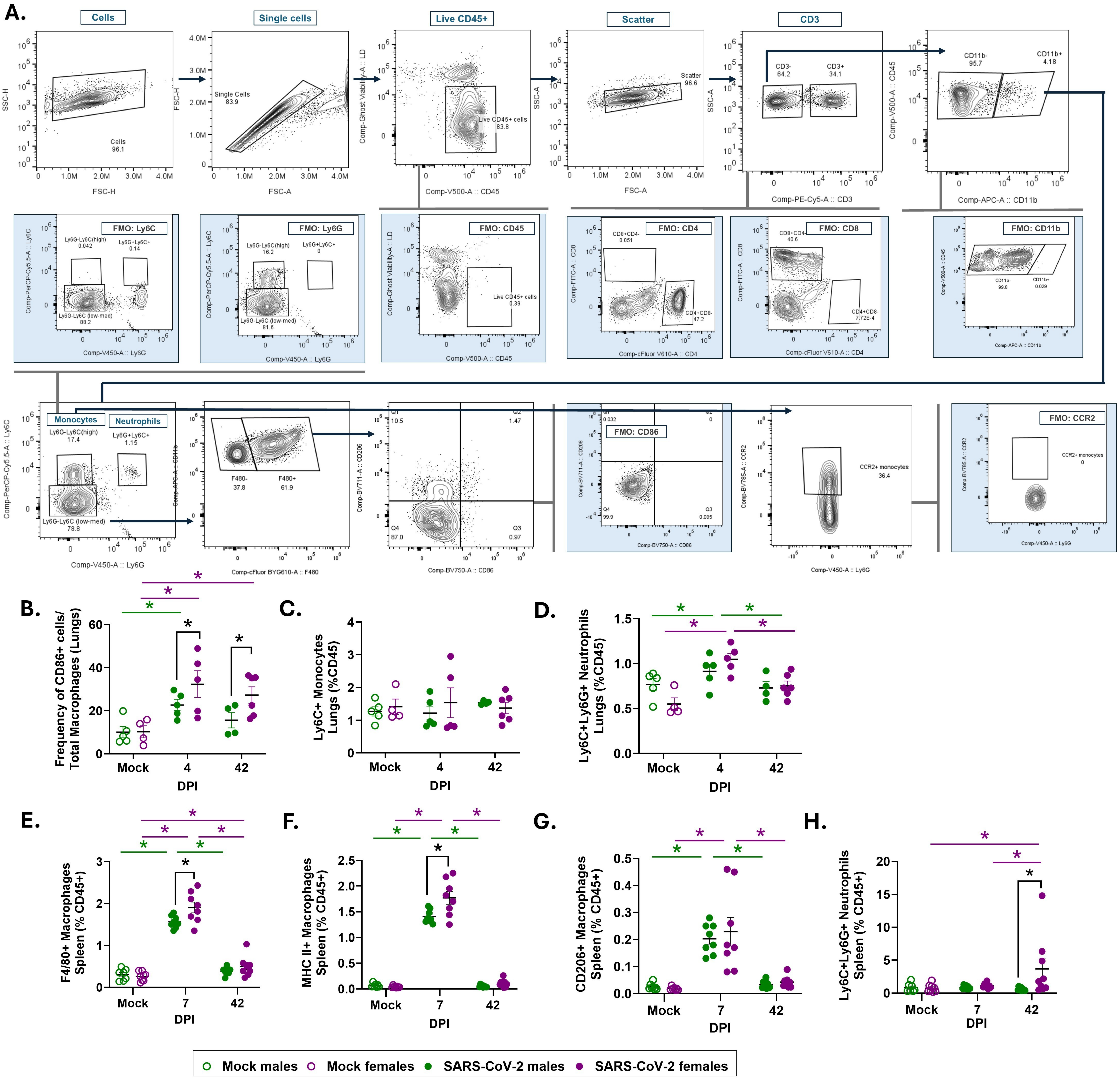

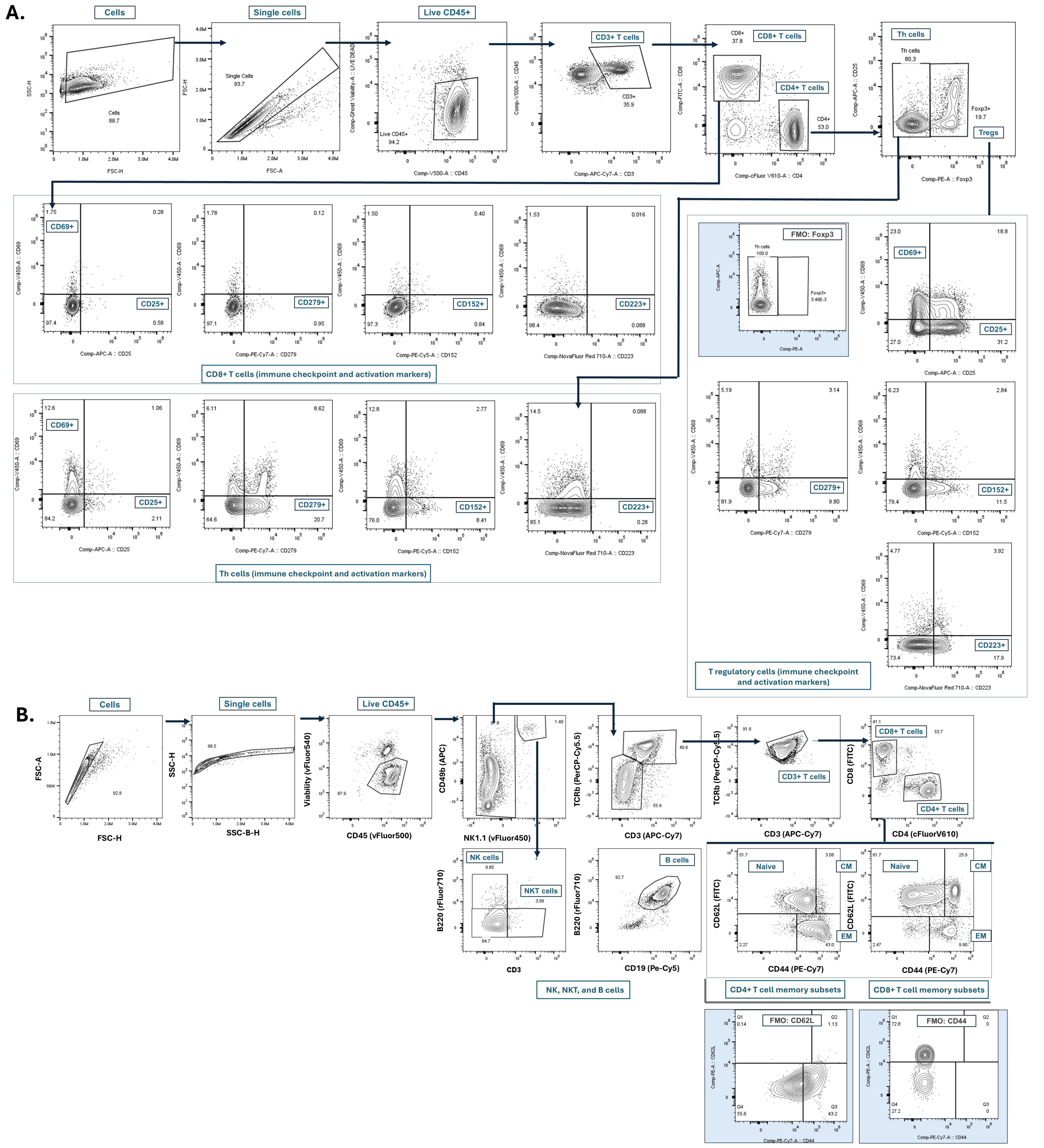

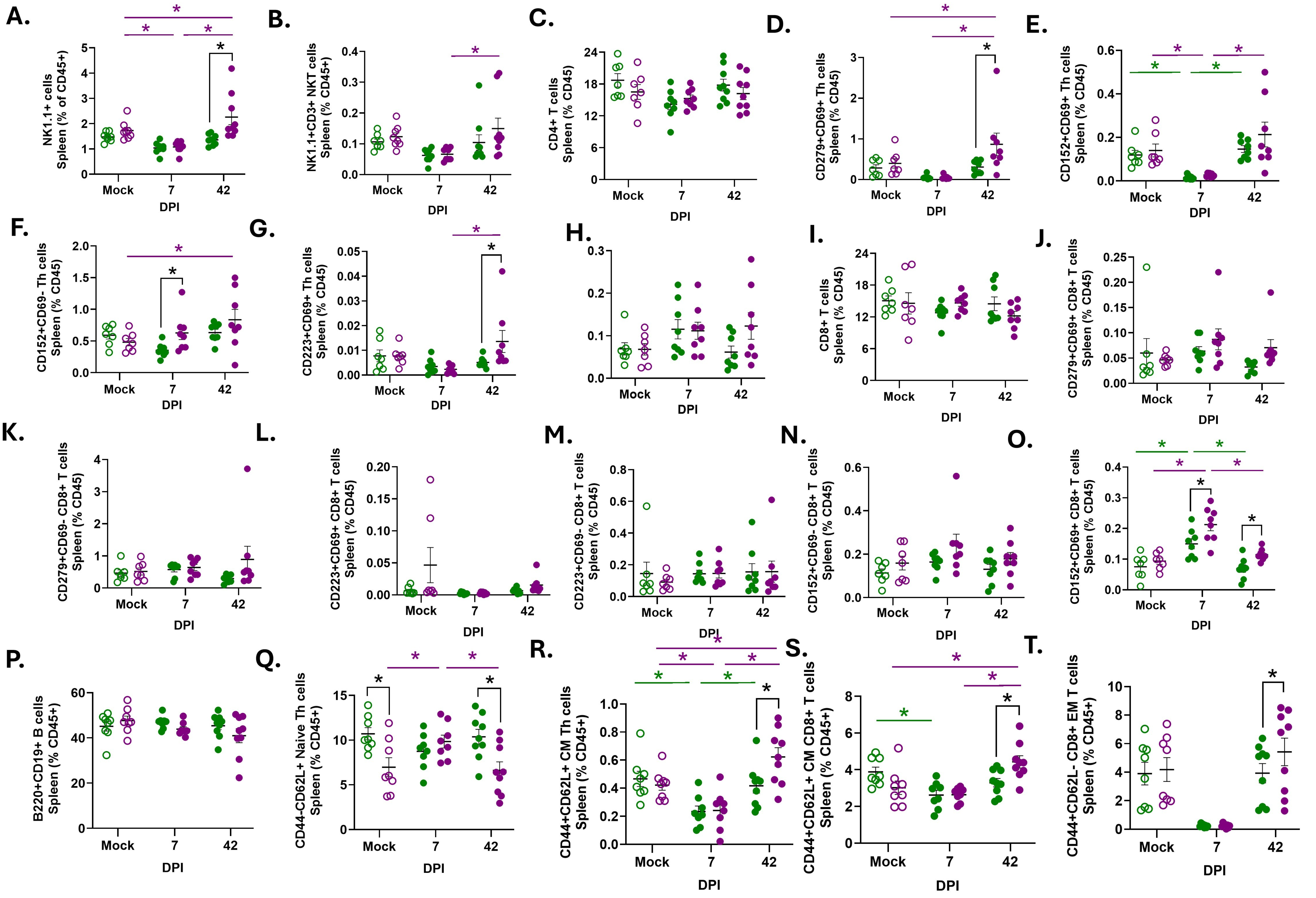

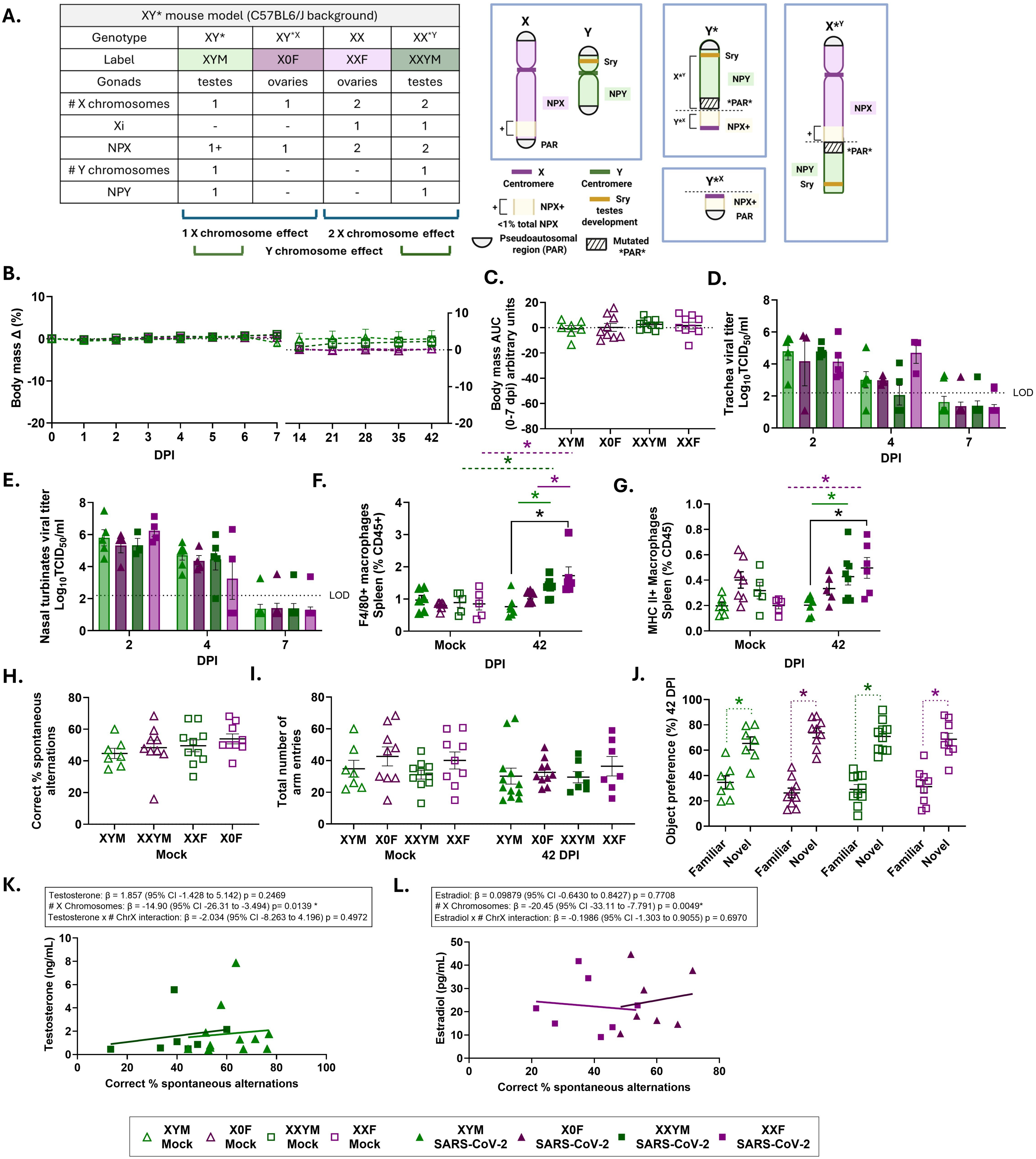

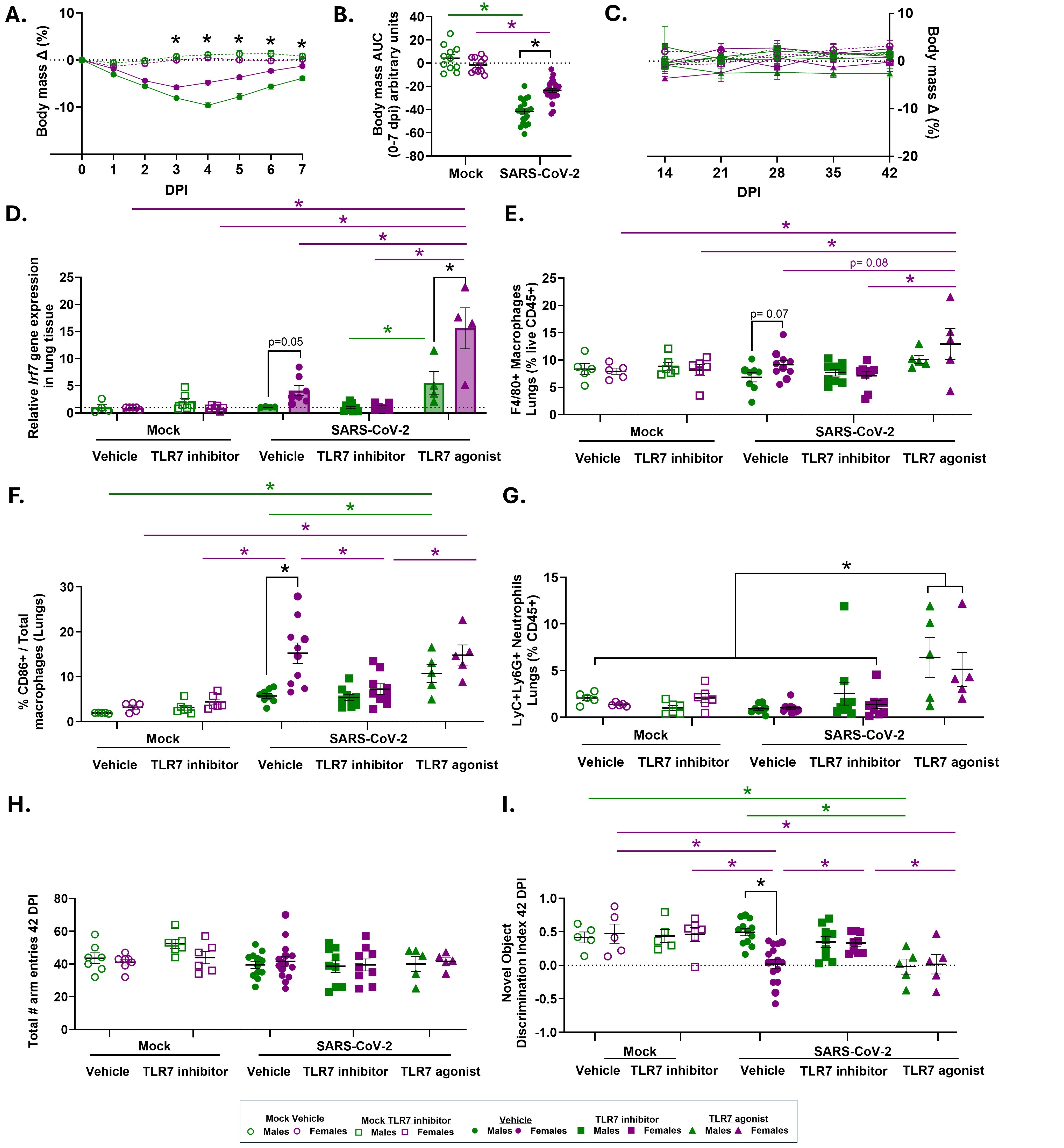
