## Supplemental Materials for "Mechanisms of sex differences in acute and long COVID sequelae in mice"

**The PDF file includes:**

**Supplemental Figure and Methods 1.** Extended behavioral graphs and methods.
**Supplemental Table 3.** Reporting animal research. list of experimental animals, strains, and
 viruses used in this study.
**Supplemental Table 4.** List of antibodies used for flow cytometry and immunofluorescence
**Supplemental Table 5.** Cell markers to identify immune cell populations.

**Extended behavioral tests in SARS-CoV-2 (B.1.621) infected mice**

.

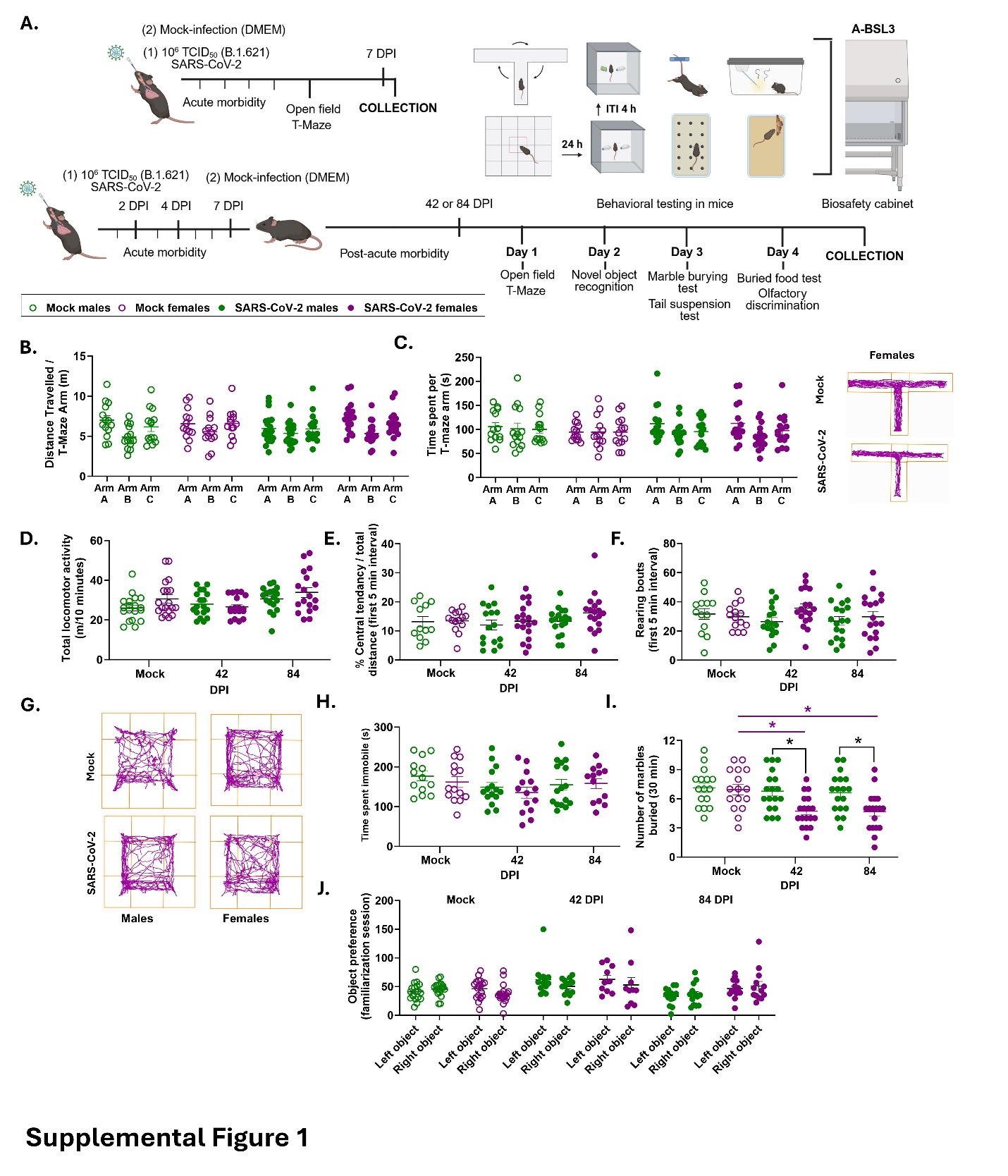

**Supplemental Figure Caption 1. Extended behavioral tests in SARS-CoV-2 (B.1.621) infected mice.** Additional behavioral measures were assessed in SARS-CoV-2 and mock-infected male and female mice at 7, 42, or 84 days post infection (dpi), using animal video tracking monitoring software (ANY-Maze) recorded in a biosafety cabinet to accommodate working in the BSL-2, or BSL-3 environment (**A**). Mock-infected control male and female mice were assessed at each timepoint and during the same time interval as SARS-CoV-2 infected mice. Because there were no significant differences among mock-infected mice based on dpi, mock-infected males and females were collapsed across time for statistical comparisons. Additional analyses and measures were conducted during the T-maze assessment (data collapsed across all timepoints since there were no within-group differences), including quantifying the total distance travelled within each arm (**B**) and the time spent in each arm (**C**) during the T-maze test; with representative locomotor track patterns for T-maze task. Locomotor activity and exploratory behaviors were assessed using the open field test (OFT), by allowing mice to freely explore an open rectangular maze during a 10-minute interval. Total distance travelled (m) was calculated at 42 and 84 dpi (**D**). Anxiety-like behaviors (i.e., central tendency (thigmotaxis), and rearing bouts) were assessed during the first 5 minutes of the OFT (**E-F)**. The total distance travelled within the center area of the maze was measured and reported as a percentage of the total distance travelled during the first 5 minute interval of the OFT (**E**). The total number of rears bouts were measured during the first 5 min interval of the OFT (**F)**, with representative activity track patterns shown (**G**). Despair behavior as an assessment of depressive-like behaviors in mice was conducted using a tail suspension test (TST) with the amount of time spent immobile while suspended quantified in seconds (s) (**H**). The buried marble test is an assessment of repetitive-compulsive like behaviors with the number of marbles buried quantified during a 30-minute test period (**I**). Object preference was assessed during the familiarization session (**J**). Data are represented as mean ± SEM from two to three independent replications (n =3-18/group). Statistical significance was determined by a two-way ANOVA followed by Tukey’s multiple-comparison test. Asterisks (*) represent significant differences (p<0.05) between groups, with black solid lines reflecting sex differences in multiple comparisons tests between males and females when a main effect of sex is observed, and colored solid lines representing either differences across timepoints post-infection, or differences within sex comparing mock and infected mice (green=males, purple=females). Open circles/bars represent mock-infected mice and closed circles/bars represent B.1.621 infected mice, with colors reflecting sex (green=males, purple=females).

**Supplemental Methods**

**Behavioral assessment**

Mice were evaluated for behavior at set timepoints following infection in the BSL-3 facilities which involves assessments within the biosafety cabinet. Behavioral assessments include measures learning and memory, including spontaneous alternation (modified T-maze assessment)^21^, and novel object recognition (NOR)^22^, measurements of locomotor and exploratory activity open field test (OFT)^74^, neuropsychiatric behaviors including the tail-suspension test (depressive-like behavior)^75^, anxiety-like behavioral assessments performed during the OFT testing interval^74^, marble burying test^76^, and assessment of olfactory response, including buried food test and the olfactory habituation/dishabituation test^77^ (**Experimental design, Extended Data 3A**). Full behavioral assessments were conducted across five consecutive days starting with the least stressful to most aversive, with the exception of olfactory assessments for the final testing day because this assessment involved an overnight fast and involved a combination of manual scoring and video monitoring using behavioral tracking software (ANYMaze) for analysis. Testing was conducted in a separate facility room from the animal holding room to minimize odor or operator interference influencing behavioral testing outcomes and were habituated for at least 30 min to the testing environment and an additional 30 min in a separate biosafety cabinet under nearly identical sound, vibration, and light exposure prior to testing. Mice were tested during the light phase due to biosafety constraints and to avoid disrupting light-dark cycles to maintain circadian rhythms.

***Open Field***

Mice were given a minimum of 1 h between behavioral tasks to re-habituate to the testing environment and biosafety cabinet. Locomotor exploratory activity was assessed during a 10 min testing interval in an OF maze constructed to fit within the biosafety cabinet. Video footage was recorded during the testing interval and the total distance travelled was analyzed using ANYmaze software. During the OFT, anxiety-like behavioral measures were assessed during the first 5 min session including measures of central tendency (thigmotaxis) by quantifying the distance traveled in the central zone compared to the total distance traveled during the first 5 min session of the OFT, and by manually scoring the number of rears performed during the first 5 min interval. After completion of the test, mice were removed from the maze and returned to their home cage. The OFT apparatus was thoroughly cleaned with Vimoba then 70% EtOH and wiped dry between trials to eliminate residual odors.

***Marble Burying Task.***
Mice were habituated to the testing room for a minimum of 30 min prior to behavioral testing. Marble burying test was performed as a secondary assessment and measure of repetitive, compulsive, or anxiety-like responses in mice. Following habituation, mice were placed individually into new, clean, and unused separate animal cages that contained a 3-4 cm thick layer of corn cob bedding. Twelve glass marbles were equally distributed into rows of 3 at equal distances from each other. Mice were given 30 min of undisturbed time to freely explore the cage, then were returned to their home cages. The number of marbles that were buried was counted during the 30 min testing interval was quantified per mouse. Marbles were considered buried with bedding if marbles were covered at least 2/3 or greater.

***Tail suspension test (TST)***
To assess depressive-like responses in mice, the TST was performed to quantify learned helplessness in a non-escapable stressful stimulation (tail-suspension). Mice were given a minimum of 1 h between behavioral tasks to re-habituate to the testing room. In a biosafety cabinet, mice were gently suspended from their tail using adhesive tape (distance from floor = ~12 in) and video recorded for a 6 min test session. Mice were returned to their home cages and the biosafety cabinet was thoroughly cleaned with Vimoba then 70% EtOH, and wiped dry between trials to eliminate residual odors. The total time mice spent immobile during the testing interval was manually scored and analyzed using ANYMAZE software. Any experimental mice that successfully climbed their tail for >30% of the total duration of time grasping their tail were excluded from analyses.

**Supplemental Table 3** | List of experimental animals, sexes, strains, and viruses in this study.

| **Experimental Animals/Strains (*Mus musculus*)** | | | |
| --- | --- | --- | --- |
| **Strain Sex** | **Source Ages at infection (weeks) Strain #** | | |
| C57BL6/J Males | Jackson Laboratory | 10, 30 | 000664 |
| Females | Jackson Laboratory | 10, 30 | 000664 |
| C57BL6/CR Males | NIA | 80 | - |
| Females | NIA | 80 | - |
| CD-1 Males | Charles Rivers | 10, 30, 80 | 027 |
| Females | Charles Rivers | 10, 30, 80 | 027 |
| XY* on C57BL/6J background | UCLA, breeding in-house | | 043694-UCD |
| Chromosomes: XY* Males |  | 30 |  |
| XXY* Males |  | 30 |  |
| XX Females |  | 30 |  |
| X0 Females |  | 30 |  |

| **Pathogen infection** | | | | |
| --- | --- | --- | --- | --- |
| **Virus** | **Lineage** | **Strain/Isolate DoseTCID_50_ Catalog #** | | |
| SARS-CoV-2 | B.1.621; Mu variant | hCoV-19/USA/MD-HP06587/2021 | 10^6^ | NR-56225 |
| SARS-CoV-2 | Mouse-adapted (Ma10) from isolate USA/WA1/2020 backbone | | 10^5^ | NR-55329 |
| **Viruses and cells:** Working stocks were generated by infecting Vero-E6-TMPRSS2 cells at a MOI of 0.01 TCID_50_ per cell infection media (DMEM) with 2.5% FBS, 100 U/mL penicillin, 100 μg/mL streptomycin, 1 mM l-glutamine, 1-mM sodium pyruvate for 72 h. | | | | |

**Supplemental Table 4** | List of antibodies used for flow cytometry and immunohistochemistry

| **Flow Cytometry (mouse)** | | | | | |
| --- | --- | --- | --- | --- | --- |
| **Cell marker** | **Fluorochrome** | | **Source** | **Concentration** | **Catalog #** |
| **Lymphoid Panel 1 (Treg panel)** | | | | | |
| CD69 | | violetFluor 450 | CytekBio | Pre-titrated | 95-K201 |
| CD45 | | violetFluor 500 | CytekBio | Pre-titrated | 95-K201 |
| Viability | | Ghost Dye violetfluor540 | CytekBio | 1:5000 | 13-0879 |
| CD4 | | cFluor V610 | CytekBio | Pre-titrated | 95-K201 |
| CD8 | | FITC | CytekBio | Pre-titrated | 95-K201 |
| Foxp3 | | PE | CytekBio | Pre-titrated | 95-K201 |
| CD152 | | PE-Cyanine5 | CytekBio | Pre-titrated | 95-K201 |
| CD279 | | PE-Cyanine7 | CytekBio | Pre-titrated | 95-K201 |
| CD25 | | APC | CytekBio | Pre-titrated | 95-K201 |
| CD223 | | redFluor 710 | CytekBio | Pre-titrated | 95-K201 |
| CD3 | | APC-Cyanine7 | CytekBio | Pre-titrated | 95-K201 |
| **Lymphoid Panel 2 (TBNK panel)** | | | | | |
| NK1.1 | | violetFluor 450 | CytekBio | Pre-titrated | 95-K101 |
| CD45 | | violetFluor 500 | CytekBio | Pre-titrated | 95-K101 |
| Viability | | Ghost dye violet540 | CytekBio | 1:5000 | 13-0879 |
| CD4 | | cFluor v610 | CytekBio | Pre-titrated | 95-K101 |
| CD8 | | FITC | CytekBio | Pre-titrated | 95-K101 |
| CD62L | | PE | CytekBio | Pre-titrated | 95-K101 |
| TCRβ | | PerCP-Cyanine5.5 | CytekBio | Pre-titrated | 95-K101 |
| CD19 | | PE-Cyanine5 | CytekBio | Pre-titrated | 95-K101 |
| CD44 | | PE-Cyanine7 | CytekBio | Pre-titrated | 95-K101 |
| CD49b | | APC | CytekBio | Pre-titrated | 95-K101 |
| CD45R (B220) | | redFluor710 | CytekBio | Pre-titrated | 95-K101 |
| CD3 | | APC-Cyanine7 | CytekBio | Pre-titrated | 95-K101 |
| **Myeloid Panel (Myeloid panel)** | | | | | |
| Ly6G | | violetFluor 450 | CytekBio | Pre-titrated | 95-K301 |
| CD45 | | violetFluor 500 | CytekBio | Pre-titrated | 95-K301 |
| Viability | | Ghost Dye violetfluor540 | CytekBio | 1:5000 | 13-0879 |
| CD8 | | FITC | CytekBio | Pre-titrated | 95-K301 |
| F4/80 | | PE | CytekBio | Pre-titrated | 95-K301 |
| Ly6C | | PerCP-Cyanine5.5 | CytekBio | Pre-titrated | 95-K301 |
| CD3 | | PE-Cyanine5 | CytekBio | Pre-titrated | 95-K301 |
| CD11c | | PE-Cyanine7 | CytekBio | Pre-titrated | 95-K301 |
| CD11b | | APC | CytekBio | Pre-titrated | 95-K301 |
| CD45R (B220) | | redFluor 710 | CytekBio | Pre-titrated | 95-K301 |
| I-A/I-E (MHC II) | | APC-Cyanine7 | CytekBio | Pre-titrated | 95-K301 |
| CD86 | | BV605 | Biolegend | Pre-titrated | 105037 |
| CD206 | | BV711 | Biolegend | 0.5 µg per 10^6^ cells | 141727 |
| CCR2 (CD192) | | BV785 | Biolegend | 1 µg per 10^6^ cells | 150621 |
| **Immunofluorescence (mouse)** | | | | | |
| **Cell marker** | |  |  | **Source Dilution** | **Catalog #** |
| Iba1 Anti-Iba1 antibody IgG rabbit monoclonal | | | | Abcam 1:1000 | ab178846 |
| P2RY12 Anti-P2RY12 antibody mouse IgG2b | | | | Biolegend 1:1000 | 848002 |
| NeuN Anti-NeuN antibody mouse polyclonal | | | | M. Sigma 1:200 | MAB377 |
| ZO-1 Anti-ZO-1 antibody rabbit monoclonal | | | | Abcam 1:200 | Ab221547 |
| DCX Anti-DCX antibody rabbit polyclonal | | | | Proteintech 1:250 | 13925-1-AP |
| DAPI Vectashield HardSet Antifade Mounting   Medium with DAPI | | | | Vector laboratories | H-1500-10 |

**Supplemental Table 5** | Cell markers to identify immune cell populations.
Table key: Underlined denotes parent population and * denotes activation or immune checkpoint receptor markers for immune cell populations.

|  | **Flow Cytometry (mouse)** | |
| --- | --- | --- |
|  | **Immune cell populations** | **Markers** |
|  | **Lymphoid Panel 1 (TBNK panel)** | |
|  | B cells | CD45+Viability-CD49b-NK1.1-CD3-B220+CD19+ |
|  | Lymphocytes | CD45+Viability-CD49b-NK1.1- |
|  | Total T cells | CD45+Viability-CD49b-NK1.1-CD3+ |
|  | CD4+ T cells | CD45+ Viability-CD49b-NK1.1-CD3+TCRβ+CD4+CD8- |
|  | Naive CD4+ T cells | CD45+ Viability-CD49b-NK1.1-CD3+TCRβ+CD8+CD4-CD62L+CD44- |
|  | Central memory (CM) CD4+ T cells | CD45+ Viability-CD49b-NK1.1-CD3+TCRβ+CD8-CD4+CD62L+CD44+ |
|  | Effector memory (EM) CD4+ T cells | CD45+ Viability-CD49b-NK1.1-CD3+TCRβ+CD8-CD4+CD62L-CD44+ |
|  | CD8+ T cells | CD45+ Viability-CD49b-NK1.1-CD3+TCRβ+CD8+CD4- |
|  | CM CD8+ T cells | CD45+ Viability-CD49b-NK1.1-CD3+TCRβ+CD8+CD4-CD62L+CD44+ |
|  | EM CD8+ T cells | CD45+ Viability-CD49b-NK1.1-CD3+TCRβ+CD8+CD4-CD62L-CD44+ |
|  | NK cells | CD45+ Viability-CD49b+NK1.1+B220+CD3- |
|  | NKT cells | CD45+ Viability-CD49b+NK1.1+B220-CD3+ |
| **Lymphoid Panel 2 (Treg panel)** | | |
| Total T cells | | CD45+ Viability-CD3+ |
| CD4+ T cells | | CD45+ Viability-CD3+CD4+CD8- |
|  | T helper (Th) CD4+ T cells | CD45+ Viability-CD3+ CD4+CD8-FoxP3- |
|  | * CD69+ | CD45+ Viability-CD3+ CD4+CD8-FoxP3-CD69+ |
|  | * CD279+CD69+ | CD45+ Viability-CD3+ CD4+CD8-FoxP3-CD69+CD279+ |
|  | * CD279+CD69- | CD45+ Viability-CD3+ CD4+CD8-FoxP3-CD69-CD279+ |
|  | * CD152+CD69+ | CD45+ Viability-CD3+ CD4+CD8-FoxP3-CD69+CD152+ |
|  | * CD152+CD69- | CD45+ Viability-CD3+ CD4+CD8-FoxP3-CD69-CD152+ |
|  | * CD223+CD69+ | CD45+ Viability-CD3+ CD4+CD8-FoxP3-CD69+CD223+ |
|  | * CD223+CD69- | CD45+ Viability-CD3+ CD4+CD8-FoxP3-CD69-CD223+ |
|  | T regulatory (Treg) | CD45+ Viability-CD3+ CD4+CD8-FoxP3+ |
|  | * CD69+ | CD45+ Viability-CD3+ CD4+CD8-FoxP3+CD69+ |
|  | * CD69+CD25+ | CD45+ Viability-CD3+ CD4+CD8-FoxP3+CD69+CD25+ |
|  | * CD25+CD69- | CD45+ Viability-CD3+ CD4+CD8-FoxP3+CD69-CD25+ |
|  | * CD279+ CD69+ | CD45+ Viability-CD3+ CD4+CD8-FoxP3+CD69+CD279+ |
|  | * CD69-CD279+ | CD45+ Viability-CD3+ CD4+CD8-FoxP3+CD69-CD279+ |
|  | CD8+ T cells | CD45+ Viability-CD3+CD4-CD8+ |
|  | * CD69+ | CD45+ Viability-CD3+CD4-CD8+CD69+ |
|  | * CD279+CD69+ | CD45+ Viability-CD3+CD4-CD8+CD69+CD279+ |
|  | * CD279+CD69- | CD45+ Viability-CD3+CD4-CD8+CD69-CD279+ |
|  | * CD152+CD69+ | CD45+ Viability-CD3+CD4-CD8+CD69+CD152+ |
|  | * CD152+CD69- | CD45+ Viability-CD3+CD4-CD8+CD69-CD152+ |
|  | * CD223+CD69+ | CD45+ Viability-CD3+CD4-CD8+CD69+CD223+ |
|  | * CD223+CD69- | CD45+ Viability-CD3+CD4-CD8+CD69-CD223+ |
| **Myeloid Panel (Myeloid panel)** | | |
| Neutrophils | | CD45+ Viability-CD3-CD11b+Ly6G+Ly6C+ |
| Monocytes | | CD45+ Viability-CD3-CD11b+Ly6G-Ly6C+(high) |
| * CCR2+ | | CD45+ Viability-CD3-CD11b+Ly6G-Ly6C+(high) CCR2+ |
| Macrophages | | CD45+ Viability-CD3-CD11b+Ly6G-Ly6C+(low-med)F4/80+ |
| * MHC II+ | | CD45+ Viability-CD3-CD11b+Ly6G-Ly6C+(low-med)F4/80+MHC II+ |
| * CD86+ | | CD45+ Viability-CD3-CD11b+Ly6G-Ly6C+(low-med) F4/80+CD86+CD206- |
| * CD206+ | | CD45+ Viability-CD3-CD11b+Ly6G-Ly6C+(low-med)F4/80+CD206+CD86- |
